# Microcephaly-Associated Genes *asp* and *Sas4* Control Chromatin Organization and Nuclear Lamina Structure in *Drosophila melanogaster*

**DOI:** 10.1101/2025.07.22.666102

**Authors:** Degisew Yinur Mengistu, Marta Marzullo, Claudia Pellacani, Marcella Marchetti, Marta Terribili, Emma Montivero Morales, Maria Patrizia Somma, Laura Ciapponi

## Abstract

Autosomal recessive primary microcephaly (MCPH) is a neurodevelopmental disorder characterized by reduced brain size and non-progressive intellectual disability. Mutations in over 30 genes have been linked to MCPH. Nearly a half of the genes identified by these mutations encode proteins involved in centrosome biogenesis or microtubule (MT) dynamics, suggesting a central role for mitotic spindle organization and division plane orientation in disease aetiology. However, it has been suggested that disruptions in spindle positioning alone are not sufficient to lead to microcephaly. Here, we investigate the contribution of the *Drosophila* orthologs of *ASPM*/*MCPH5* (*asp*) and *CENPJ/MCPH6* (*Sas4*) to nuclear architecture, chromatin organization, and genome stability. We show that loss of either Sas4 or Asp leads to aberrant microtubule architecture, mislocalization of the LINC complex, and deformation of the nuclear lamina. These defects are accompanied by reduced levels of both lamin and HP1α and impaired centromere clustering in interphase cells. Sas4 and asp mutants also exhibit a global reduction in heterochromatin-associated histone marks (H3K9me2/3 and H3K27me3) and increased levels of the euchromatin-associated mark H3K4me3. Remarkably, treatment with Methylstat, a demethylase inhibitor, reduced nuclear invaginations by partially restoring H3K9me3 levels. Additionally, Sas4 or Asp depletion leads to DNA damage, increased sensitivity to genotoxic stress, and delayed DNA repair.

Together, these findings reveal a previously underappreciated role for Asp and Sas4 in preserving nuclear architecture and chromatin integrity, offering new insight into the pathogenesis of MCPH.

**GRAPHICAL ABSTRACT:** 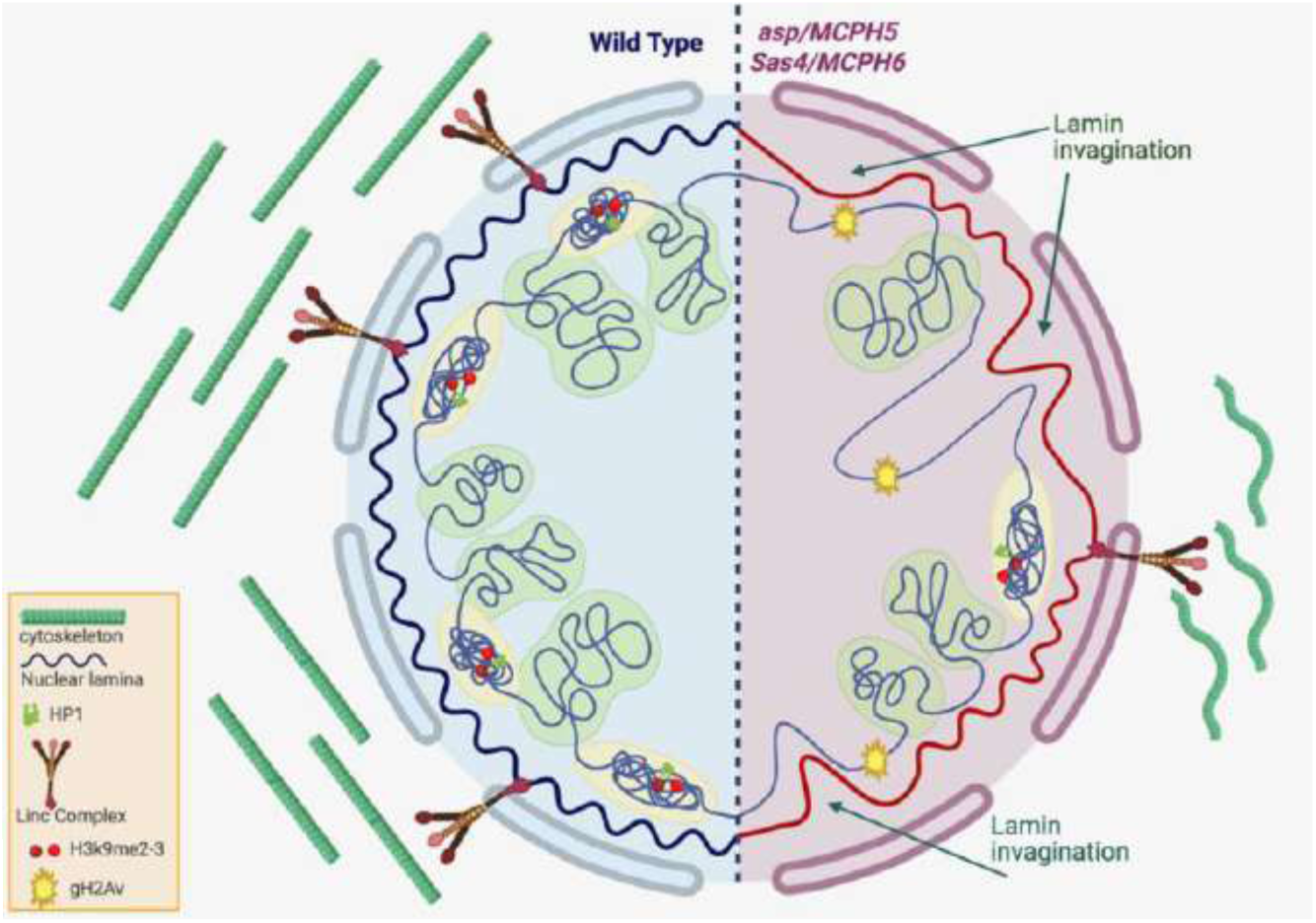

## INTRODUCTION

Autosomal Recessive Primary Microcephaly (MCPH) is a neurodevelopmental disorder present at birth and characterized by reduction of occipitofrontal head circumference (OFC) by at least three standard deviations from the mean [1–5]. MCPH results from neurogenesis defects leading to a reduced number of neurons. To date, MCPH has been linked to recessive mutations in 30 genes [6], which are expressed in all proliferating cell types that will eventually form the cerebral cortex. These genes encode proteins involved in many cellular processes that are essential for proper brain development, including the balance between proliferation and differentiation, the timely migration of neurons, and the maintenance of genome stability [7,8]. Specifically, 12 of the 30 MCPH genes encode centrosomal components and/or regulators of MT dynamics [9]. Centrosome defects can lead to the disruption of mitotic spindle positioning which may result in an imbalance between symmetric and asymmetric division in neural stem cells (NSCs) [10-12). This imbalance would reduce the proliferation of neural progenitors and/or cause premature neuronal differentiation, which eventually leads to the generation of a small-sized brain.

However, recent evidence indicates that altered spindle positioning during neurogenesis is not sufficient to directly induce significant changes in cell fate [8,13,14]. Thus, mitotic defects do not fully account for the complexity of the MCPH etiology.

DNA damage accumulation has been observed in many MCPH models, consistent with the role of several MCPH proteins in DNA repair and genome stability [15]. However, the pool of neural progenitor cells could also be significantly depleted by modifying the fate of proliferating neural stem cells and inducing their premature differentiation. This might be due to the premature activation of lineage-specific genes and the concomitant silencing of stem cell genes. The establishment of such a transcriptional program must be tightly coordinated with cell division and might be accompanied by changes in chromatin organization. Recently, a great emphasis has been given to the impact of post translational histone modifications on the spatiotemporal gene expression programs during neurogenesis [16]. The N-terminal histone tail is subjected to many modifications (including lysine acetylation, lysine and arginine methylation, serine/threonine phosphorylation, etc.) that regulate the accessibility of the associated DNA regulatory elements.

Specifically, methylation of histone H3 on lysines 4, 36, 79 (H3K4, H3K36 and H3K79) is generally associated with poised or active gene transcription, whereas methylation of histone H3 on lysine 9, 20, 27 (H3K9, H3K20 and H3K27) is a hallmark of gene silencing in the heterochromatic regions. Among those histone repressive marks, H3K27me3 is crucial for the regulation of the progression of neural stem cell (NSC) lineages [17].

Emerging evidence suggests that mechanical forces from the cytoskeleton contribute to the regulation of chromatin organization and nuclear function during neurogenesis. Interestingly, recent work suggests that chromatin organization is affected by forces exerted by the cytoskeleton on the nucleus [18,19]. The connection between the nucleus and the cytoplasm is mediated by the MT-actin cytoskeleton and components of the nuclear membrane, such as the LINC complex and the lamin nucleoskeleton. The lamin meshwork is essential to establish the 3D architecture of the genome by anchoring heterochromatic regions to the nuclear envelope [20]. The resulting chromatin compartmentalisation is required for the regulation of many nuclear processes, such as transcriptional activation and repression and DNA damage response and chromatin dynamics during the cell cycle. Lamin dysfunction causes a heterogeneous group of diseases known as laminopathies [21]. A common feature of laminopathyc cells is the presence of misshaped nuclei and heterochromatin loss (needs a ref). The correlation between altered MT cytoskeleton, heterochromatin remodelling and lamin dysfunction has been demonstrated in a *Drosophila* tauopathy model where the aberrant cytoskeletal-nucleoskeletal interaction, leads to the heterochromatin relaxation, loss of H3K9me2 and HP1α promoting neurodegeneration [22,23].

The *ASPM/MCPH5* (Abnormal SPindle-like Microcephaly-associated) gene, the human ortholog of the *Drosophila asp* gene, has been found mutated in over 40 % of MCPH cases. Asp/ASPM proteins localize to the MT minus ends [24] and play a crucial role in the regulation of mitotic spindle assembly and orientation [25]. ASPM is a component of a protein complex required for centriole biogenesis and activity [26].This complex is formed by WDR62, CEP63 and CENPJ, all of which are proteins associated with microcephaly [12]. Although Asp/ASPM was initially thought to function exclusively during mitosis, recent studies have revealed its involvement in interphase processes such as DNA repair and protein ubiquitination [11, 27, 28]. The centromere-associated protein J (*CENPJ/MCPH6*) gene, the human ortholog of *Drosophila Sas4* (Spindle Assembly Abnormal 4), encodes a centrosomal protein involved in centriole duplication and centrosome assembly. CENPJ/Sas4 mediates the interaction between the pericentriolar material (PCM) scaffold and the centriole during centrosome biogenesis by providing a platform termed “extended surface-like” [29, 30]. Depletion of mouse CENPJ protein leads to loss of centrosomes and defective brain development [31,32].

In this study, we investigated whether cytoskeletal dynamics and nuclear morphology are disrupted in *Drosophila* models of MCPH resulting from the depletion of either the *Sas4* or *asp* gene. We found that these genes are critical for the maintenance of chromatin architecture and genome integrity during *Drosophila* neurogenesis. Deletion of these genes, besides the well-known effects on spindle structure and function, cause reduction in heterochromatin marks, lamin invaginations and hypersensitivity to DNA damage. Altogether, our data suggest that Sas4 and Asp are essential for maintaining a balance between proliferation and differentiation by affecting not only the mitotic process but also the architecture and morphology of the nucleus.

## RESULTS

### Cytoskeleton defects in *asp* and *Sas4* mutant brain cells

To explore the connection between cytoskeleton and nuclear architecture, we examined the microtubule (MT) network in interphase neuroblast (NB) cells lacking either Sas4 or Asp in both mutants (*asp^t25^*, *Sas4^s2214^*) and RNAi-brains (RNAi experiments are shown in the supplementary figures S4-S5 and S1 Data). Neuroblasts were initially identified by staining with a Deadpan antibody, which recognizes a transcription factor specific to stem cells. Subsequently, we measured the nuclear diameter of Deadpan-positive cells and restricted our analysis to those with a diameter greater than 10 μm.

In wild-type (wt) interphase cells, the centrosome is easily identifiable as a discrete focus located close to the nucleus and serves as the primary site of microtubule (MT) nucleation [33]. From this site, long MTs radiate outward to form a well-organized network that spans the entire cell. In *Sas4* mutant NBs, however, the centrosome is not detectable, and MTs are instead arranged in bundles restricted to the cell periphery, rather than forming an integrated network (Figure 1A). This phenotype resembles that observed in neurons following *CENPJ* silencing [34]. In *asp* mutant NBs, the MT network appears disorganized and consists of short, fragmented MTs (Figure 1A). The linescan intensity profile analyses provide a clear visualization of the difference in MT organization patterns. While the fluorescence intensity peaks in wt NBs are indicative of well-organized MT bundles, in Sas4 mutant NBs appear to lack MT bundles except for those located beneath the nuclear membrane. In asp mutant NBs, the low level of MT bundle organization is reflected by the high fluorescence baseline and the absence of prominent intensity peaks in the corresponding linescan. The connection between the cytoskeleton in the cytoplasm and the nuclear envelope is maintained by the LINC (Linker of Nucleoskeleton and Cytoskeleton) complex. To determine if Asp and Sas4 are required for proper LINC distribution, we stained larval brain NBs from wild-type and mutant flies with an antibody against Klarsicht (Klar), a LINC complex component. We found that in wild type NBs, Klar was mainly located at the nuclear envelope, while *Sas4* and *asp* mutant cells showed Klar dispersed throughout the nucleus (Figures 1B and C). These results prompted us to determine whether the loss of Sas4 and Asp alters the morphology of the lamin-based nucleoskeleton. We performed co-immunostaining both for tubulin and LaminDm0, the *Drosophila* ortholog of human Lamin B. Altered morphology of lamin meshwork was observed in both *Sas-4* and *asp* mutant neuroblasts and, noteworthy, MT bundles frequently localized near these lamin invaginations (Figure 1D-F). This abnormal nuclear morphology phenotype was associated with significantly reduced levels of Lamin protein, as detected in wt and mutant brain extracts by Western blotting (Figure 1G). Collectively, our data reveal that Sas4 and Asp are important for maintaining the proper connection between the cytoskeleton and the nuclear envelope.

**Figure 1:**
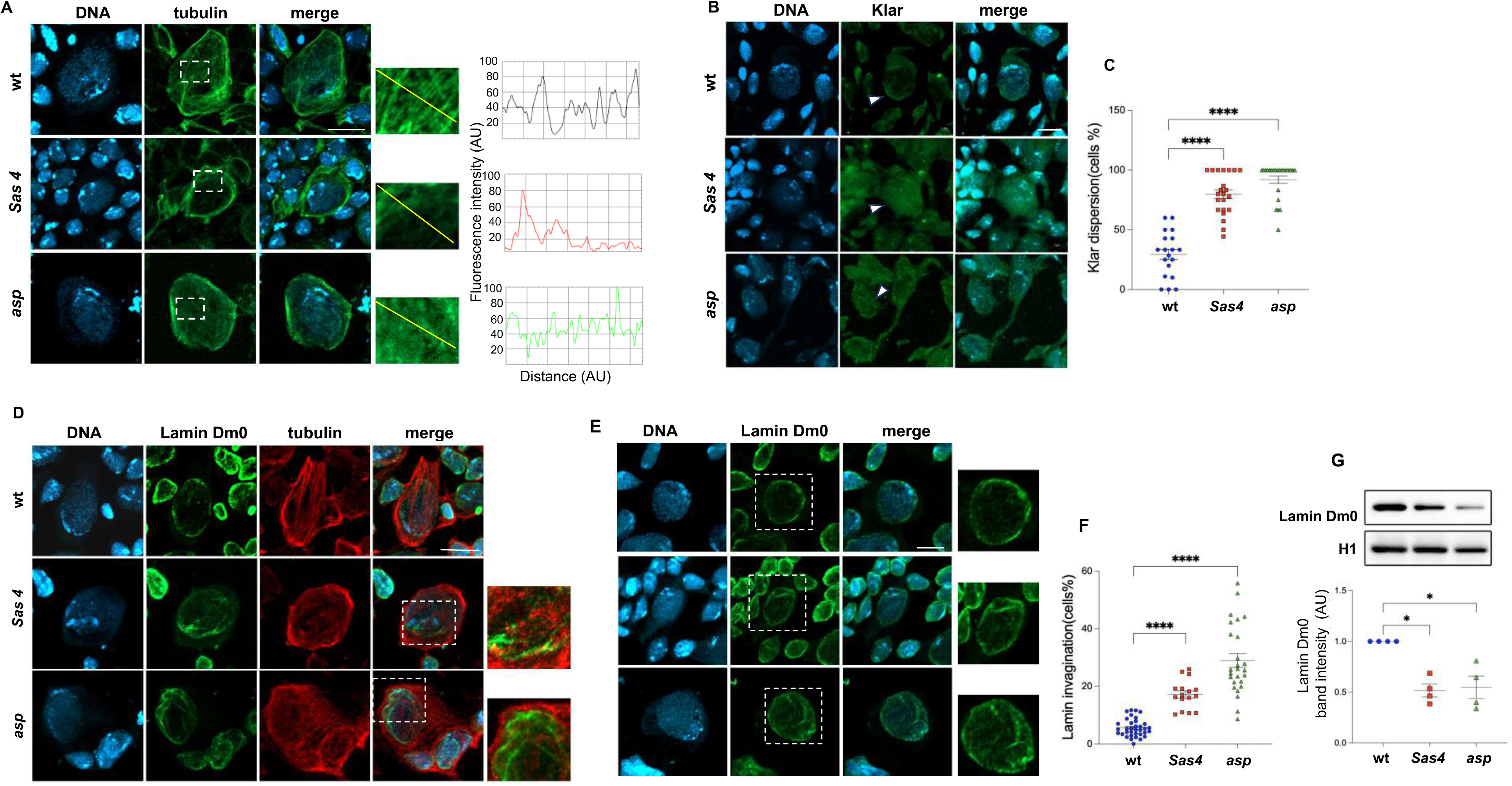
Cytoskeleton defects in *asp* and *Sas4* mutant brain cells. **A)** Neuroblast cells from third instar brain squashes of wild type (wt) and *Sas4* or *asp* mutants immunostained with anti-tubulin antibody (green) and DAPI (blue, DNA). Dotted lines indicate the cell portion shown at higher magnification on the right, with corresponding tubulin intensity profiles (black for wild type, red for *Sas4* and green for *asp* mutants) across the highlighted segment. B) Immunostaining of wt and *Sas4* or *asp* larval brain cells using the anti-Klarsicht (Klar) antibody (green) and DAPI (blue, DNA). Note that Klar is localized around the nuclear envelope (arrowhead) in wt neuroblasts, whereas it is diffused into the nucleus (arrowheads) in mutant neuroblasts. C) Graphical representation of the percentage of cells showing Klar dispersion in wt brain cells (blue rounds), and in *Sas4* (red square) or *asp* (green triangles) mutants. Each dot represents the percentage of cells per 63x microscope field in 3 wt and 3 mutant brains (n≥20). D) Neuroblast cells from third instar brain squashes of wt and *Sas4* or *asp* mutants immunostained with anti-LaminDM0 (green) and anti-tubulin (red) antibodies. DAPI is in blue (DNA). Dotted lines indicate the cells selected for the magnification shown on the right. E) Neuroblast cells from third instar brain squashes of wt and *Sas4* or *asp* mutants immunostained with anti-LaminDM0 antibody (green) and DAPI (blue, DNA). Dotted lines indicate the cells selected for the magnification shown on the right. F) Graphical representation of the percentage of cells showing invaginations of the nuclear envelope in *Sas4* (red rectangles) and *asp* (green triangles) mutant and wt (blue circles) brain cells. Each dot represents the score of cells per 63x microscope field in 3 wt and 3 mutant brains (n≥20). G) Representative immunoblots on protein extracts from wt, *Sas4* or *asp* larval brains labelled with anti-LaminDM0 antibody, with the corresponding band quantification normalized on the loading control (H1) in at least three independent experiments. AU, arbitrary unit. Error bars represent SEM. P = p-value calculated using unpaired t test. *p < 0.05; **p < 0.01; ***p < 0.001; ****p < 0.0001. Scale bar = 10 μm.

### Loss of Asp and Sas4 affects heterochromatin organization and leads to nuclear envelope dysfunction in larval brain cells

To assess whether the Lamin defects in *Sas4* and *asp* mutants lead to altered heterochromatin positioning, we analysed the localization of the centromere-specific H3 variant (CENP-A; Centromere identifier, Cid in flies) in early prophase, when the spindle microtubules start to be nucleated. It is known that Cid signals are clustered near the apical centrosome in wild type NBs [35]. Early prophase in *asp* mutant NBs were identified not only by the size but also by the initial assembly of asters and the presence of Cid signal doublets belonging to sister chromatids. In contrast, in *Sas4* mutants that lack centrosomes and asters, prophase NBs were identified solely based on the size and the presence of Cid signal doublets. We found that centromere clustering was reduced in both *asp* and *Sas4* mutant cells, as indicated by an increased ratio of the area occupied by Cid signals to total nuclear area (Figure 2A and B).

**Figure 2:**
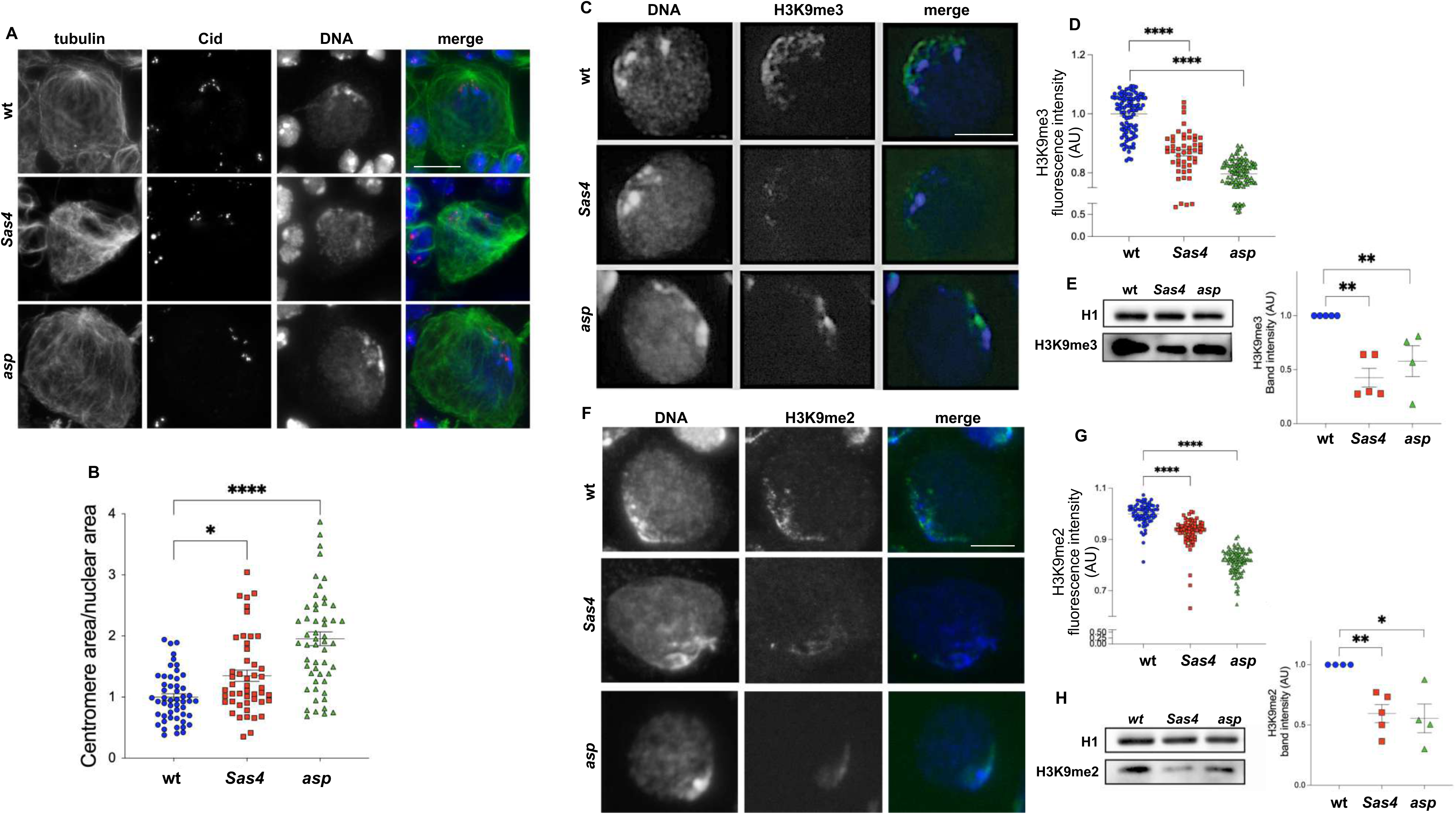
Asp and Sas4 are required for centromere clustering and H3K9me2-3 positioning. A) Neuroblast cells from third instar brain squashes of wt and *Sas4* or *asp* mutant brains immunostained with the anti-centromere specific Histone-3 variant marker, Cid, and anti-tubulin antibodies. In merged images Cid is in red, tubulin is in green and DNA in blue (DAPI). Note that the Cid signals are clustered and localized close to the apical centrosome in wild type neuroblast prophase, while in S*as4* and *asp* mutant prophases the centromeres were dispersed. B) Graphical representation of the centromere area expressed as the ratio between the area occupied by Cid signals and the total nuclear area in brain cells of wt (blue rounds) and *Sas4* (red square) or *asp* (green triangles) mutants. Each dot represents a single nucleus from at least 3 brains (n ≥ 50). C) Neuroblast cells from third instar larval brain squashes of wt and *Sas4* or *asp* mutants stained with anti*-* H3K9me3 antibody and DAPI (DNA). In merged images H3K9me3 is in green and DNA in blue. D) Quantification of H3K9me3 fluorescence intensity per cell as in (C), each dot represents a single cell from at least 3 brains (n≥50). E) Representative western blots showing decreased levels of H3K9me3 in *Drosophila* larval brain extracts from *Sas4* or *asp* mutants, compared to wt with the corresponding quantification of relative band intensity on loading control (H1), in at least three independent experiments. F) Neuroblast cells from third instar larvae brain squashes of wt and *Sas4* or *asp* mutants stained with anti-H3K9me2 antibody and DAPI (DNA). In merged images H3K9me2 is in green and DNA in blue. (G) Quantification of H3K9me2 fluorescence intensity per cell as in (F), each dot represents a single cell from at least three brains (n ≥ 50). H) Representative western blots showing decreased levels of H3K9me2 in *Drosophila* larval brain extracts of *Sas4* or *asp* mutants compared to wt, with the corresponding quantification of relative band intensity on loading control (H1) in at least three independent experiments. AU, arbitrary unit. Error bars represent SEM. P = p-value calculated using unpaired t test. *p < 0.05; **p < 0.01; ***p < 0.001; ****p < 0.0001. Scale bar = 10 μm.

Since in human cells lamin invaginations are frequently associated with heterochromatin relaxation [23,36], we examined the distribution of H3K9me2 and H3K9me3, two key epigenetic marks of gene silencing that guide the peripheral positioning of silent chromatin. Immunostaining of larval brain cells from *asp* and *Sas4* mutants and fluorescence quantification showed a significant reduction in methylation levels compared to controls (Figure 2C and D, 2F and G), an observation confirmed by Western blotting analysis of brain protein extracts (Figure 2E and H). Notably, in mutant brains co-stained with LaminDm0 and H3K9me3, the nuclear envelope invaginations were consistently associated with a marked reduction of these heterochromatic marks (Figure 3A and B). Importantly, all cell types in Sas4 and asp mutant brains exhibited a substantial decrease in H3K9me2 and H3K9me3, as well as lamin invaginations.

**Figure 3:**
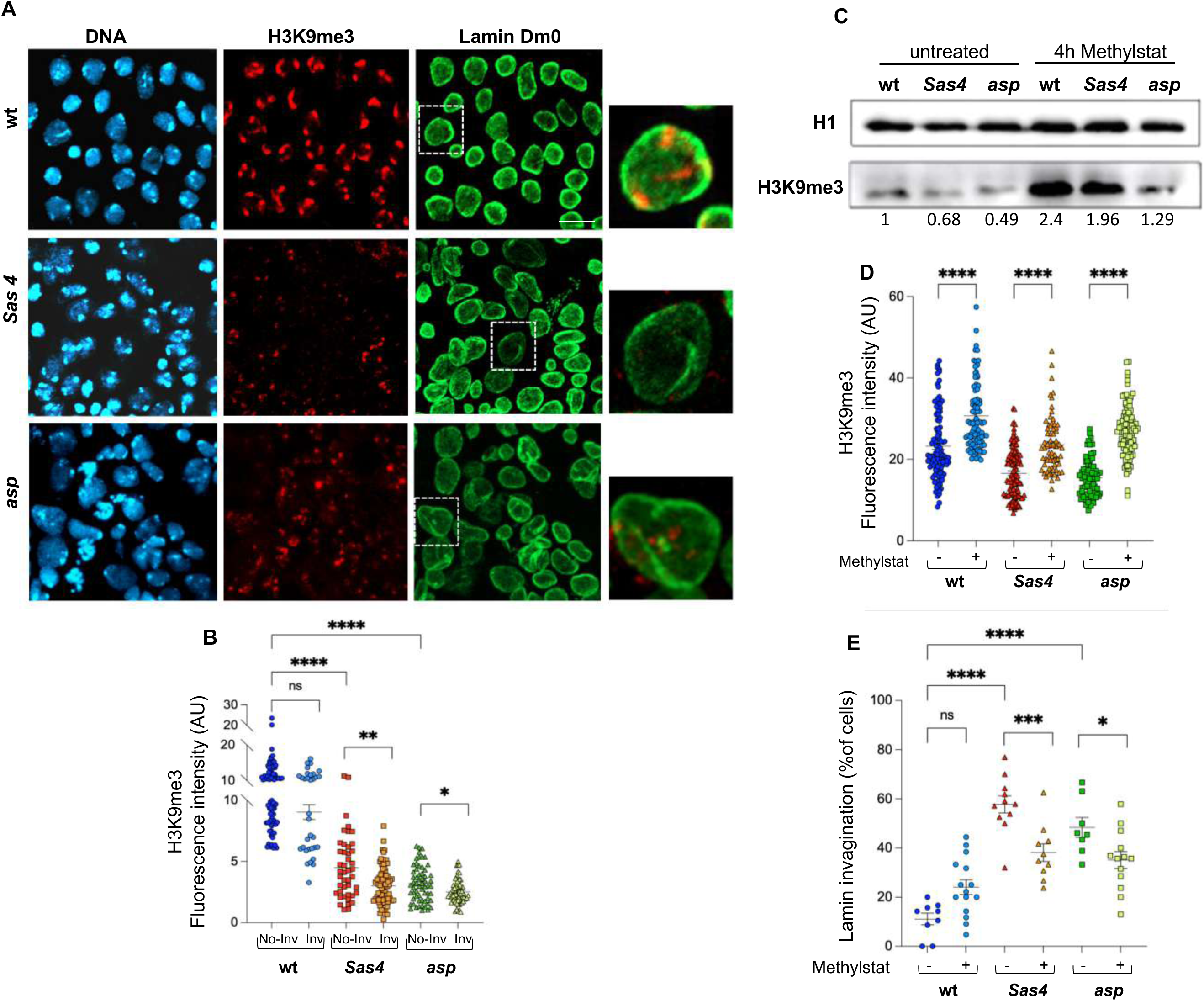
Lamin invaginations and lamin level reduction in *Sas4* and *asp* mutant brain cells are dependent on H3K9me3 levels. A) Third instar brain squashes of wt and *Sas4* or *asp* mutant immunostained with anti-LaminDM0 (green) and H3K9me3 (red) antibodies, and DAPI (blue, DNA). Dotted lines indicate the cells selected for the magnification shown on the right. Scale bar = 10 μm. B) Fluorescence intensity quantification of H3K9me3 signal in cells with normal nuclear envelope or in cells with lamin invaginations in *Sas4* mutant (red and orange squares), *asp* mutant (green and light green triangles) and wt (blue and light blue rounds) showing a significant correlation between lamin invaginations and H3K9me3 reduction. Each dot represents a single cell (n ≥ 30). C) Representative immunoblots on protein extracts from wt, *Sas4* or *asp* larval untreated brains or upon 4h treatment with 3μM Methylstat, labelled with anti-LaminDM0, below the corresponding band quantification normalized on the loading control (H1), is indicated. D) Fluorescence intensity quantification of H3K9me3 in *Sas4* mutant (red and orange squares) and *asp* mutant (green and light green triangles) and wt (blue and light blue rounds) untreated brain cells treated or upon 4h treatment with 3μM Methylstat. Each dot represents a single cell from at least 3 wt and 3 mutant brains (n ≥ 50). E) Graphical representation of the percentage of cells showing invaginations of the nuclear envelope in *Sas4* (red and orange squares) or *asp* mutants (green and light green triangles) and wt (blue and light blue rounds) brain cells without treatment or after 4h treatment with 3μM Methylstat. Each dot represents the score of cells per 63x microscope field in at least 3 wt and 3 mutant brains (n ≥ 8). Arbitrary Unit (AU). Error bars represent SEM. P = p-value calculated using unpaired t test. *p < 0.05; *p < 0.05, ***p < 0.001 ****p < 0.0001.

Consistent with this result, *ex vivo* treatment of mutant larval brains with methylstat, a demethylase inhibitor [37], partially restored H3K9me3 level (Figure 3C and D) and significantly reduced the Sas4- or Asp-dependent nuclear invagination phenotype (Figure 3E). This suggests that restoring proper H3K9me3 levels through methylstat treatment helps to re-establish normal nuclear architecture by maintaining heterochromatin in developing brain cells.

Consistent with these results, we also found significantly reduced total levels of heterochromatin protein 1α (HP1α), a specific component of heterochromatic regions (Figure 4). HP1 binds to H3K9Me2/3, localizes to heterochromatin and its binding to methylated H3K9 plays an important role in the establishment and maintenance of 3D genome organization during development [38–40]. HP1α is enriched in heterochromatic foci, which are readily visible in control brains stained with an HP1α antibody (Figure 4A). Interestingly, there is no corresponding accumulation of HP1α at the residual H3K9me3 foci detected in *Sas4* and *asp* mutant brain cells (Figure 4A). Quantification of HP1α fluorescence revealed reduced signal intensity in mutant larval brain cells compared to controls (Figure 4B), which was associated with reduced HP1α levels as detected by Western blot (Figure 4C).

**Figure 4:**
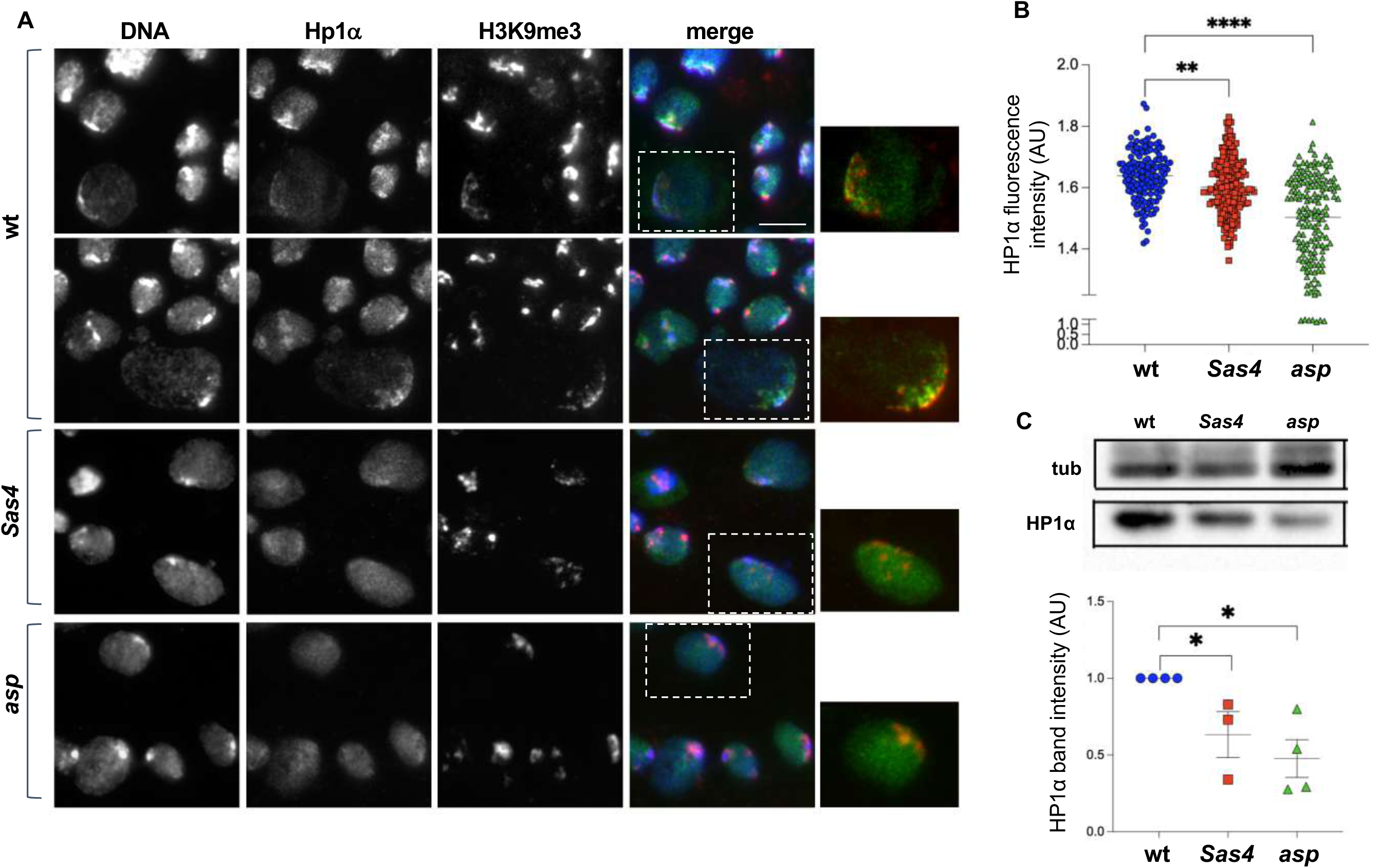
HP1α is reduced and delocalized in the nucleus of *Sas4* and *asp* mutant cells. A) Immunostaining of wt and *Sas4* or *asp* mutant third instar larval brains stained with anti-HP1α and H3K9me3 antibodies and DAPI (DNA). In merged images HP1α is in green, H3K9me3 in red and DNA is in blue. Note that HP1α accumulates in specific regions of the nucleus in wt cells, whereas in mutant cells, HP1α is distributed throughout the nucleus. Scale bar = 10 μm. B) Graphical representation of fluorescence intensity quantification of HP1α in brain cells of wt (blue rounds), and *Sas4* (red square) or *asp* (green triangles) mutants. Each dot represents a single cell (n ≥ 140). C) Representative immunoblots for HP1α on protein extracts from wt, *Sas4* or *asp* larval brains with the corresponding band quantification normalized on the loading control (tubulin) in at least three independent experiments. AU, Arbitrary Unit. Error bars represent SEM. P = p-value calculated unpaired t test. *p < 0.05; *p < 0.05, ***p < 0.001 ****p < 0.0001.

We next assessed the state of facultative heterochromatin, which is critical for silencing lineage-specific genes during *Drosophila* larval brain development [17]. To this end, we analysed H3K27me3 levels in *Sas4* and *asp* mutant larval brains. Immunostaining revealed reduced H3K27me3 signal intensity in mutant neuroblasts compared to wild-type cells (5A and B). However, western blot analysis of whole brain extracts showed no significant difference in overall H3K27me3 levels among wild-type, *asp*, and *Sas4* mutants (Figure 5C). This discrepancy likely reflects the fact that H3K27me3 levels remain high in differentiated cells in both mutant and control brains, while the reduction is specific to neuroblasts (Figure 5A, B, and D). Interestingly, analysis of the euchromatin mark H3K4me3 revealed an inverse pattern, with increased levels in mutant neuroblasts that were not detected by Western blotting (Figure S1).

**Figure 5:**
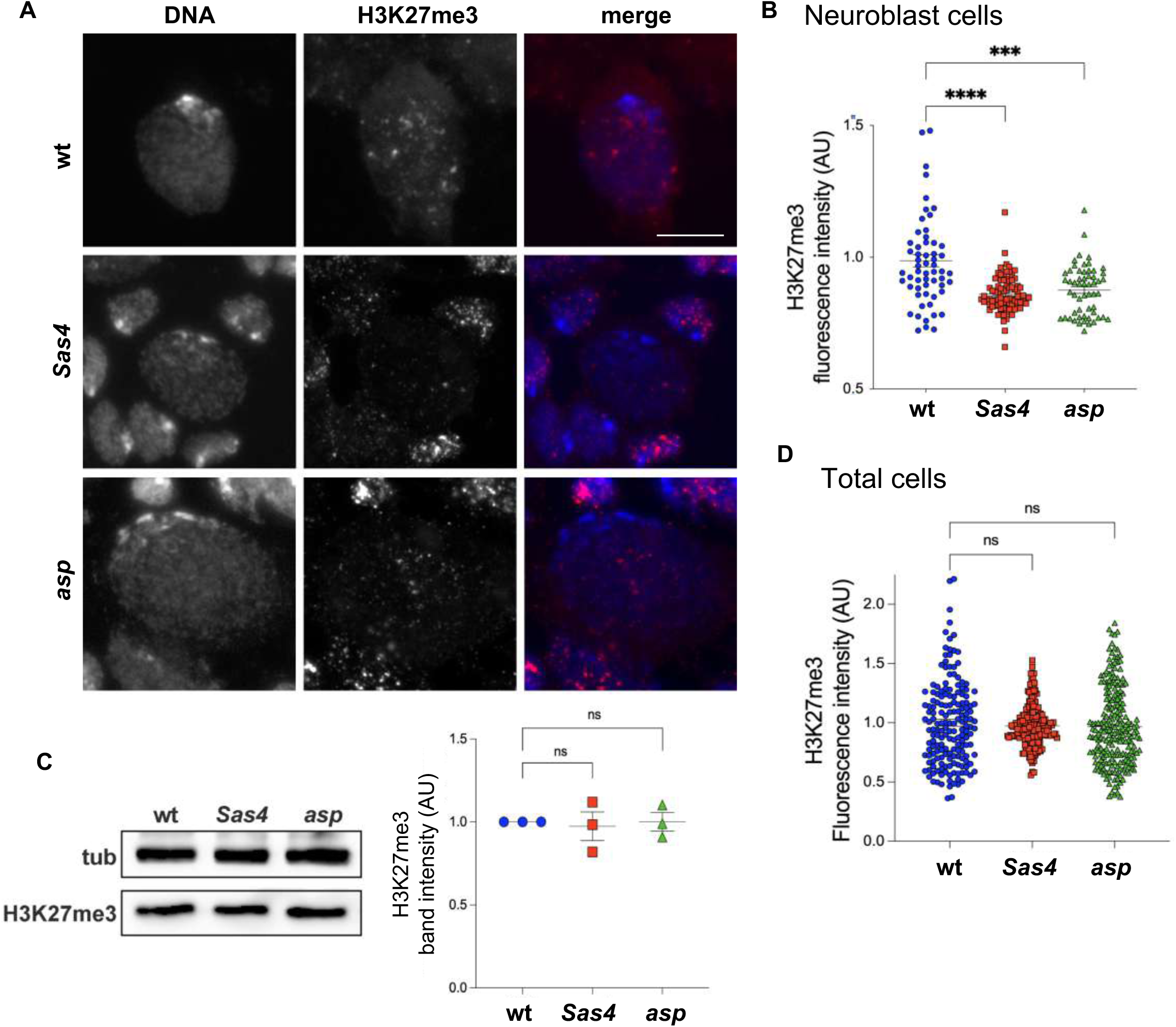
The facultative heterochromatin mark, H3K27me3, is reduced in *Sas4* and *asp* mutant neuroblasts. A) Immunolocalization of H3K27me3 on brain squashes from third instar larvae of wt and *Sas4* or *asp* mutants. In the merged panels, H3K27me3 is in red and DNA in blue (DAPI). Scale bar = 10 μm. B) Fluorescence intensity quantification of H3K27me3 in neuroblasts labelled as in (A), showing a decrease of H3K27me3 in both *Sas4* (red square) and *asp* (green triangles) mutant cells, compared to wt (blue rounds; n ≥ 50). C) Representative western blots showing similar levels of H3K27me3 in larval brain extracts of *Sas4* and *asp* compared to wt and the corresponding band quantification normalized on the loading control (tubulin) in at least three independent experiments. D) Fluorescence intensity quantification of H3K27me3 in all larval brain cells/field; each dot represents a single cell (n ≥ 150). Arbitrary Unit (AU). Error bars represent SEM. P = p-value calculated using unpaired t test; ns, not significant; *p < 0.05; *p < 0.05, ***p < 0.001 ****p < 0.0001. Scale bar = 10 μm.

Taken together, these data indicate that loss of either Sas4 or Asp reduces Lamin levels and disrupts lamin nucleoskeleton architecture, potentially impairing nuclear envelope integrity. This structural alteration may in turn contribute to the altered heterochromatin positioning and relaxation, indicating a functional role for Sas4 and Asp in the maintenance of nuclear architecture and heterochromatin in brain development.

### Polytene chromosome banding patterns are altered in *asp* and *Sas4* mutant salivary glands

Polytene chromosomes from *Drosophila* salivary glands are characterized by a chromocenter that contains under-replicated heterochromatin and bands of intercalary heterochromatin also in euchromatic arms. This characteristic offers a unique model for studying the distribution and arrangement of chromatin marks on interphase chromosomes [41,42]. To explore the pattern and dynamics of chromatin marks in *asp* and *Sas4* mutants, we performed immunostaining of polytene chromosomes from squashed third-instar salivary glands. Fluorescence quantification analysis confirmed a near-complete loss of H3K9me3/me2 signals, especially in the chromocenter, which is normally a strong signal-rich region due to its heterochromatic nature. This loss suggests a substantial reorganization or decondensation of heterochromatin in the mutant context (Figure 6A and B). A significant reduction in the repressive H3K27me3 mark was also observed (Figure 6C and D), along with a mild but significant increase in H3K4me3 euchromatic mark (Figure 6F and G).

**Figure 6.**
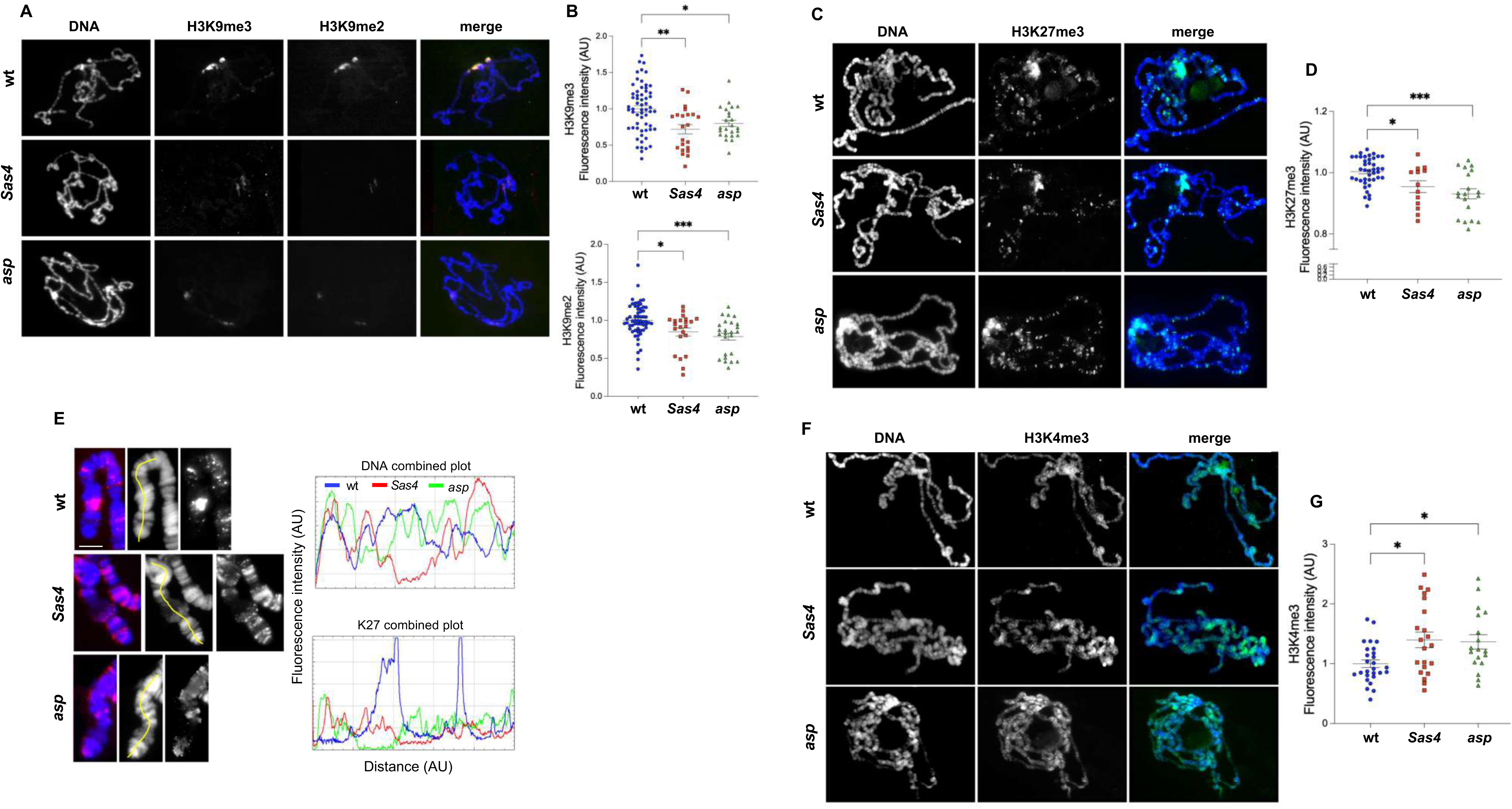
H3K9me2/3 and H3K27me3 marks are reduced and altered in *Sas4* and *asp* mutant polytene chromosomes, while the euchromatin mark H3K4me3 is increased. A) Immunolocalization of H3K9me3 and me2 on wt and *Sas4* or *asp* mutant polytene chromosomes. In merged images H3K9me3 is in green, H3K9me3 in red and DNA (DAPI) is in blue. B) Fluorescence intensity quantification of H3K9me2 and me3 signals showing a significant decrease of both heterochromatic marks in mutants compared to wt. Each dot represents a single polytene chromosome (n ≥ 25). C) Immunolocalization of H3K27me3 on wt and *Sas4* or *asp* mutant polytene chromosomes. In merged images H3K27me3 is in green and DNA in blue (DAPI). D) Fluorescence intensity quantification of H3K27me3 signals showing a significant decrease of this heterochromatic mark in mutants compared to wt. Each dot represents a single polytene chromosome (n ≥ 15). E) Examples of the X chromosome extremities stained with anti-H3K27me3 (red) and DNA (blue, DAPI) in wt and *Sas4* or *asp* mutant polytene chromosomes with corresponding intensity profiles showing an altered pattern of DNA and H3K27me3 signals in both *Sas4* (red line) and *asp* (green line) mutants compared to wt (blue line); y-axis: fluorescence intensity; x-axis: distance from the tip of the chromosome. Scale bar = 5 μm. F) Immunolocalization of H3K4me3 on wt and *Sas4* or *asp* mutant polytene chromosomes. In merged images H3K4me3 is in green and DNA in blue (DAPI). G) Graphical data quantification of fluorescence intensity of H3K4me3 as that shown in (F). Each dot represents a single polytene chromosome (n ≥ 15). AU, arbitrary unit. Error bars represent SEM. P = p-value calculated using unpaired t test. *p < 0.05; **p < 0.01; ***p < 0.001; ****p < 0.0001.

Interestingly, the band signal profiles of both DNA and H3K27me3 were altered in polytene chromosomes from *asp* and *Sas4* mutants, suggesting an extensive reorganization of intercalary heterochromatin, and possibly euchromatin as well (Figure 6E and Figure S2). These changes in the DNA banding pattern further support our hypothesis that Asp and Sas4 loss disrupts chromatin organization, potentially altering gene expression programs during brain development

### Sas4 and Asp are required for genome integrity maintenance and DNA damage response

One of the pathological hallmarks of microcephaly is genome instability caused mainly by persistent double-strand breaks (DBS) during development [43,44]. Previous work has shown that the human orthologues of *Sas4* and *asp*, [28,31], are not only required for proper spindle assembly and function but also to prevent DNA damage. Here, we investigated whether inactivation of these genes affects DNA repair and leads to DNA damage. We analysed the levels of phosphorylated H2Av (γH2Av), a well-known marker of DNA damage, both by immunostaining and western blotting. Cytological analysis of γH2Av-positive cells revealed a significant increase in mutant larval brains compared to wild type (Figures 7A and B), in line with the elevated γH2Av levels observed by Western blot (Figure 7C). These findings indicate that mutations in *Sas4* and *asp* lead to increased endogenous DNA damage, highlighting their crucial role in maintaining genome stability during *Drosophila* brain development.

**Figure 7:**
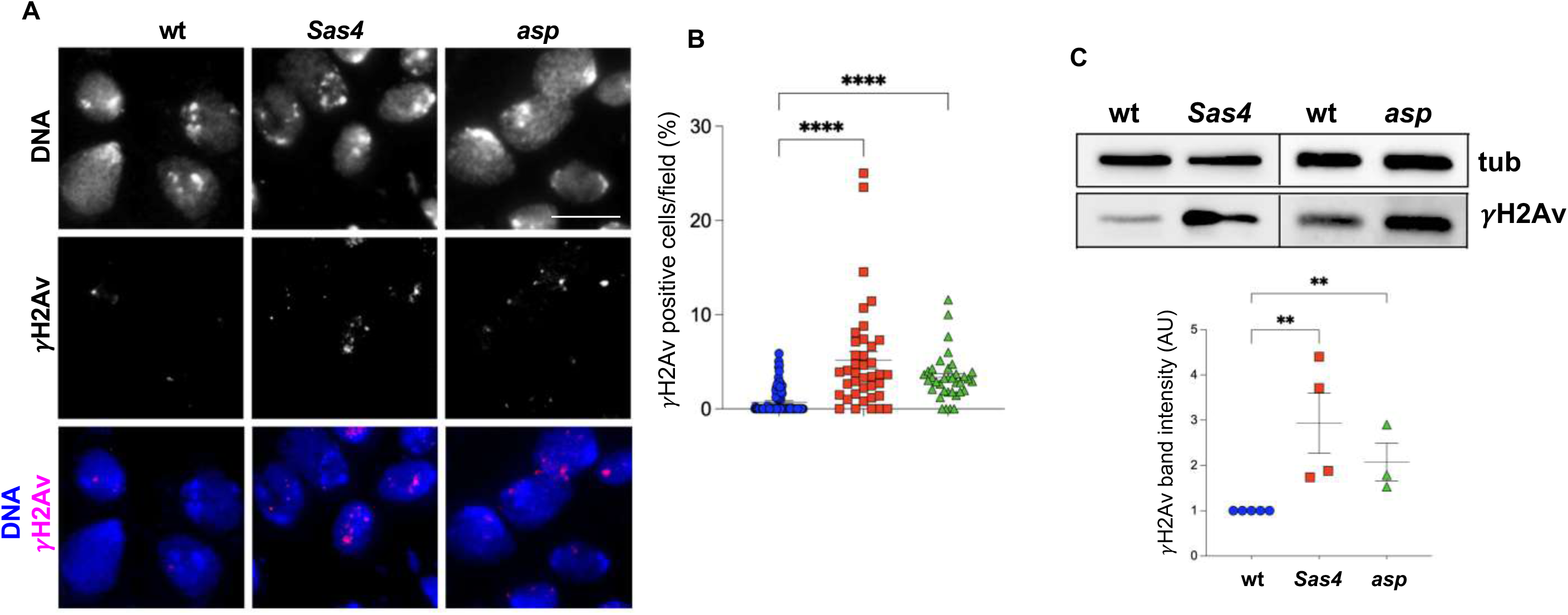
Loss of Sas4 and Asp causes increased levels of γH2Av. A) Immunostaining of wt and *Sas4* or *asp* mutant third instar larval brain cells stained with anti-γH2Av antibody and DAPI (DNA). B) Graphical representation of the percentage of γH2Av positive cells per field, as shown in (A). n≥35 for 3 brains. Scale bar = 10 μm. C) Representative immunoblots on brain extracts from wt and *Sas4* or *asp* mutant labelled using anti-γH2Av antibody with the corresponding band quantification normalized on the loading control (Tubulin). Error bars represent SEM. P = p-value calculated using unpaired t test. *p < 0.05; **p < 0.01; ***p < 0.001; ****p < 0.0001.

To understand if silencing of *asp* and *Sas4* also causes hypersensitivity to induced DNA damage, we treated mutant brains with X-rays. The larval brains were exposed to 3 or 10 Gy, dissected at different times, from 5 min to 2 hours post-irradiation (PI), and then analysed to determine the kinetics of DNA damage repair. Western blot analysis showed a significant increase in γH2Av level compared to the wild type at all recovery times. Particularly, at 2 hours PI with 3 Gy, the level of γH2Av was significantly decreased in wild type while it remained high in the mutants (Figure 8A). Similar results were obtained by using 10 Gy or after 2h hydroxyurea (2mM) treatment (Figure S3). Consistent with the Western blot results, immunofluorescence analysis revealed a significant increase in the number of γH2Av foci in the mutant cells compared to wild type, both at 5 minutes and 2 hours after treatment (Figures 8B-D). These results indicate that the DSB repair ability in the *Sas4* and *asp* mutant cells was impaired and significantly delayed, resulting in hypersensitivity to X-ray induced DNA damage.

**Figure 8:**
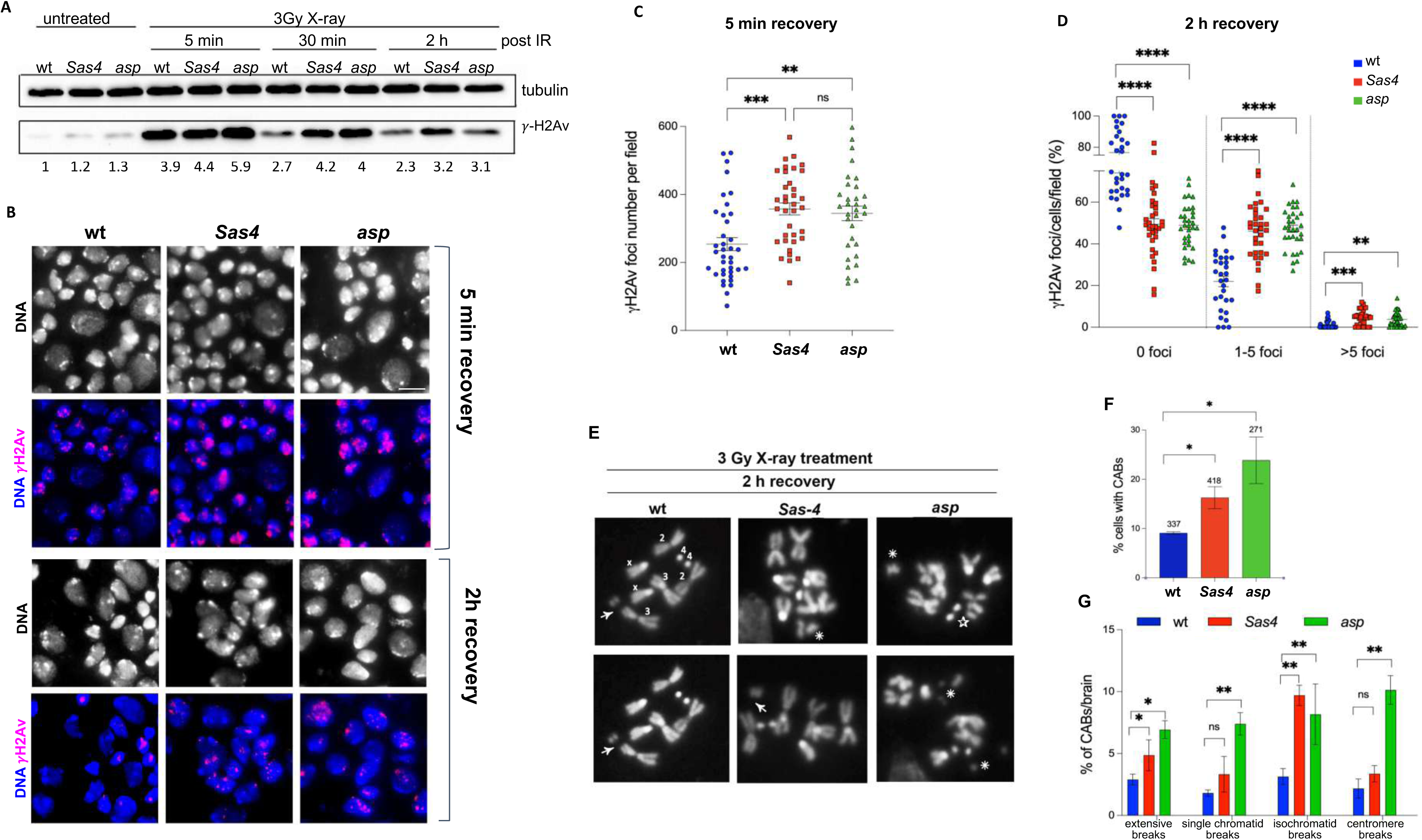
Loss of *Sas4* or *asp* induces delayed DNA damage recovery and enhanced radiosensitivity. A) γH2Av levels in wt, *Sas4* and *asp* larval brain extracts after irradiation with 3Gy and dissection after 5 min, 30 min and 2 hours post irradiation (PI). Below the corresponding band quantification normalized on the loading control (Tubulin), is indicated. B) *Drosophila* brain squashes from wt, *Sas4* and *asp* larvae, irradiated with 3Gy and stained with anti-γH2Av antibody at 5 min or 2 hours PI. Scale bar = 10 μm. C) Quantification of the γH2Av foci per field (n≥30, from at least 3 brains) at 5 min PI time, as shown in (B). C) Graphical representation of the percentage of γH2Av positive cells per field (n ≥ 30) with 0, 1-5 or >5 foci per cell, in brain cells of wt (blue rounds), *Sas4* (red square) and *asp* (green triangles) mutants of at least 3 brains, at 2 hours PI time, as shown in (B). E) Examples of larval brains metaphase spreads from wt and *Sas4* or *asp* mutants at 2 hours PI. Note: the presence of iso-chromatid breaks (arrow), extensive DNA fragments (asterisks) and chromatid deletions (star) in mutant cells. F) Graphical representation of chromosome aberrations (CABs) frequencies after 3Gy treatment in wt, and *Sas4* or *asp* mutant brain cells. Numbers above the columns represent the number of total cells analysed on at least 3 different larval brains. G) Graphical representation of the percentage of different types of CABs after 3Gy treatment observed in wt, *Sas4* or *asp* mutant larval brain metaphase spreads. AU, arbitrary unit. Error bars represent SEM. P = p-value calculated using unpaired t test. *p < 0.05; **p < 0.01; ***p < 0.001; ****p < 0.0001.

Because double-strand DNA breaks generate both γH2Av foci and chromosomal aberrations (CABs) [45], we examined CABs frequency in *Sas4* and *asp* larval brain preparations. Third instar larvae were irradiated with 3 Gray and then colchicine-treated metaphase analysis was performed at 2 hours PI. As expected, in Sas4 or Asp depleted brain cells, we found a significant increase in metaphase cells with chromosomal aberrations and extensive chromosome fragmentation compared to wild type (Figure 8E-G).

Taken together, these data demonstrate that loss of either Sas4 and Asp causes hypersensitivity to DNA damage and a failure to resolve DSBs in larval brains, suggesting that these proteins are involved in the DNA damage response and play a key role in maintaining genome integrity.

## DISCUSSION

### The role of Sas4 and Asp in establishing and maintaining nuclear architecture

In this study, we investigated the role of *Sas4* and *asp*, orthologs of human *CENPJ* and *ASPM*, in molecular pathways underlying MCPH pathogenesis. A hallmark of MCPH is a reduction in neuron number during development, often caused by imbalances in the tightly regulated processes of cell proliferation, differentiation, and apoptosis. Within this context, we have shown that Sas4 and Asp function not only in classical mitotic roles, such as centrosome function and spindle orientation, but also in the organization of the interphase cytoskeleton, as shown by the altered MT architecture observed in *Sas4* and *asp* mutant brain cells.

The cytoskeleton acts as a structural integrator of nuclear shape and positioning. Disruption of its organization leads to deformation of the nuclear lamina, affecting chromatin organization and genome stability. Studies in *Drosophila* embryos have shown that MT polymerization produces pushing forces that affect nuclear envelope dynamics and promote chromatin mobility [46]. Altered nuclear architecture, characterized by nuclear envelope invaginations, reduced Lamin levels, and mislocalization of envelope proteins, has been linked to both neurodevelopmental and neurodegenerative disorders [47]. Notably, studies in *Drosophila* tauopathy models have shown that lamin dysfunction can drive chromatin decompaction and DNA damage [22,23].

Consistent with this, we found that Sas4 or Asp loss leads to reduced lamin, relaxed heterochromatin, and frequent nuclear envelope invaginations. These features, along with impaired Klarsicht distribution, reflect broad nuclear architectural defects. Moreover, reduced centromere clustering and altered abundance and localization of HP1α suggest that Sas4 and Asp are essential for proper heterochromatin compartmentalization. The interaction between H3K9 methylation and HP1α is critical for organizing 3D genome architecture; its disruption destabilizes pericentric heterochromatin. Studies in murine embryonic stem cells have shown that HP1 loss compromises self-renewal and pluripotency, partly through reduced H3K9 methylation [48].

In line with these findings, we observed decreased levels of H3K9me2, H3K9me3, and H3K27me3 in *Sas4* and *asp* mutant brains, alongside increased H3K4me3, a mark of active chromatin, suggesting a shift toward transcriptional activation. Interestingly, total H3K27me3 levels were unchanged in whole-brain extracts, indicating cell-type specificity: the mark was reduced in neuroblasts but remained stable in differentiated cells. This points to differential regulation of repressive chromatin during development.

Further support comes from polytene chromosome analysis in salivary glands, where we observed altered banding patterns and reduced intensity of H3K9me2/3 and H3K27me3. These changes imply disrupted gene expression programs crucial for specific developmental stages and cell-type specification [17,49].

Altogether, our data suggest that loss of Sas4 or Asp disrupts heterochromatin organization and 3D genome architecture, contributing to impaired gene regulation, defective cell cycle progression, and ultimately, developmental abnormalities. Notably, H3K9 methylation plays a key role in maintaining cell identity by preventing inappropriate gene reprogramming and supports neuronal plasticity by silencing non-lineage genes during development [50,51].

### The role of Sas4 and Asp in DNA/chromosome stability and integrity

Several MCPH genes encode proteins involved in the DNA damage response [15]. For instance, MCPH1 promotes homologous recombination repair [52,53], while ASPM (MCPH5) facilitates the repair of double strand breaks by stabilizing BRCA1 [54]. Deletion of *KNL1* (*MCPH4*) in vivo leads to DNA damage on missegregated chromosomes and triggers rapid apoptosis [55]. Similarly, *CITK* deficiency results in double-strand breaks, increased sensitivity to ionizing radiation, and cell cycle arrest [56]. Additionally, CENPJ, which is associated with Seckel syndrome, has been linked to elevated DNA damage and apoptosis in a corresponding mouse model [31].

We found that the loss of Sas4 or Asp leads to a significant increase in γH2Av foci and delayed recovery from DNA damage following exposure to genotoxic stress. Several DNA damage response (DDR) proteins, including ATR, ATM, 53BP1, and DNA-PKcs, are known to localize to centrosomes, and microtubules are thought to act as tracks for transporting these proteins to the nucleus [57, 58]. Furthermore, at least for ASPM, a direct interaction with known factors involved in DNA damage repair has been demonstrated [54,59,60]. Our data suggest that Sas4 and Asp may also contribute to genome integrity through a distinct function. Given the roles of Sas4 and Asp in centrosome biogenesis and MT organization, it is plausible that they contribute to DDR regulation through these cytoskeletal functions. In addition, recent studies have revealed that cytoplasmic MTs, in concert with motor proteins and the LINC complex, facilitate the movement of DNA damage sites within the nucleus, thereby promoting repair by relocating lesions to specialized, repair-conducive nuclear domains [61–63]. In this context, Sas4 and Asp may support these processes by maintaining MT integrity and enabling damage site mobility. Another key factor influencing DNA repair efficiency is the local chromatin environment. While chromatin relaxation is often required to allow repair machinery access to DNA lesions, decondensed chromatin is more susceptible to DNA damage compared to compacted heterochromatin [64–68]. Thus, disturbances of chromatin organization not only impair repair but may also increase the accumulation of DNA damage. Importantly, defects in nuclear architecture, including abnormal nuclear morphology, increased DNA damage, and altered chromatin compaction, are common features of various pathologies, such as premature aging syndromes and cancers, many of which involve mutations in nuclear envelope components [69].

Taken together, these observations support the hypothesis that the impaired DNA damage recovery observed in *Sas4* and *asp* mutant cells may result from nuclear envelope dysfunction and heterochromatin loss, linking centrosome and cytoskeletal defects to compromised genome integrity. Remarkably, our findings emphasize the potential role of the MT cytoskeleton in the epigenetic dysregulation that underlies the aetiology of microcephaly.

## CONCLUSIONS

In conclusion, emerging evidence highlights the crucial role of a proper coordination between the microtubule/actin cytoskeleton, the LINC complex, the nuclear envelope, and heterochromatin in maintaining nuclear and chromatin dynamics to regulate genome organization, expression, and stability [18,19, 70]. Disruption of any component within this axis can lead to aberrant gene expression by altering the spatial organization of chromosomes [71]. Indeed, numerous diseases, including cancer and neurodegenerative disorders, are associated with defective microtubule networks, abnormal nuclear morphology, and genomic instability.

Considering our findings on *Sas4* and *asp* mutant cells, and considering that their human orthologs, *CENPJ* and *ASPM*, are classified as *MCPH* genes, it is plausible that the observed reduction in heterochromatin marks may lead to aberrant gene expression. This misregulation could prematurely activate lineage-specific transcriptional programs during neurogenesis, resulting in impaired or immature neuronal differentiation.

Recently, transcriptional analysis of *asp* mutant *Drosophila* larval brains revealed a global downregulation of temporal transcription factors and upregulation of immune-related genes [72]. However, the study could not resolve transcriptional signatures at the single-cell level. To date, the only available scRNA-seq data comes from an *ASPM* ferret model, which showed preserved transcriptional programs but altered proportions of neural progenitors [73]. While further studies are needed, these findings support our hypothesis that *Sas4* and *asp* regulate spatiotemporal gene expression during brain development.

This model may also extend to explain the molecular mechanisms underlying microcephaly caused by other *MCPH*-associated mutations. Indeed, at least 20 of the 30 known *MCPH* genes encode factors that are essential for the assembly and function of key cellular structures, including the centrosome, microtubules, chromatin, and the nuclear envelope, which all work together to preserve proper chromatin architecture. Taken together, our findings emphasize the potential role of the MT cytoskeleton in the epigenetic dysregulation that underlies the aetiology of microcephaly.

## MATERIALS AND EXPERIMENTAL METHODS

### *Drosophila* strains and rearing conditions

*Drosophila* stocks were maintained at 18° C for long term, whilst the stocks used in the experiments were maintained and bred at 25° C. For RNAi we used *tubulin-GAL4* to induce expression of the UAS-RNAi construct and the fly crosses were incubated at 29° C to increase the efficiency of RNAi. Fly stocks used in this study are listed in table 1.

**Table 1:** *D. melanogaster* genetic resource table.

| Stock Designation | Source or reference | Identifiers | Additional information |
| --- | --- | --- | --- |
| *tubulin-GAL4* | Bloomington | 5138 | *y[1] w[\*]; P{w[+mC]=tubP-GAL4}LL7/TM3, Sb[1] Ser[1]* |
| *Sas4<sup>s2214</sup>* | Bloomington | 12119 | *w1118; P{lacW}Sas-4s2214/TM3, Sb1* |
| *asp<sup>1</sup>* | Bloomington | 1972 | *In(3R)Ubx[7LL]ats[R], asp[1] ats[1] p[p]/TM6B, Tb[1] ca[1]; y[1]/Dp(1;Y)y[+]* |
| *asp<sup>t25</sup>* | [26] | - | - |
| *UAS asp<sup>RNAi</sup>* | Bloomington | 28741 | *y[1] v[1]; P{y[+t7.7] v[+t1.8]=TRiP.JF03169}attP2* |
| *UAS Sas4<sup>RNAi</sup>* | Bloomington | 35049 | *y[1] sc[\*] v[1] sev[21]; P{y[+t7.7]v[+t1.8]=TRiP.HMS01463}attP2* |
| *W<sup>1118</sup>* | Bloomington | - | - |

### X-ray and HU treatment

X-ray treatment was carried out using MHF PLUS (Portable X-ray unit-Gilardoni). Third instar *Drosophila* larvae were irradiated with 3 and 10 Gy, and dissected after 5 min, 30 min, 2 hours and 6 hours. Hydroxyurea treatment was performed by incubating larval brains for 30 min in 2mM HU and then in tissue culture medium for 2 hours. Brains were then collected for subsequent analyses.

### Methylstat treatment

The third instar larvae were dissected, and the brains were collected and treated with 3 μM Methylstat in Schneider’s medium for 4 hours. Subsequently, after lysis, the brain protein extracts from each sample were subjected to Western blot analysis. For immunohistochemistry, the brains were fixed with 3.7% formaldehyde and followed next steps as described below.

#### Immunohistochemistry

The samples were incubated in 3.7% formaldehyde for 20 min, transferred for 30 s in 45% acetic acid for 30 sec, and fixed in 60% acetic acid for 2 min. Successively, the brains were squashed and frozen in liquid nitrogen. After the removal of the coverslips, the slides were placed in cold ethanol for 15 min, rinsed twice for 10 min in 0.1%TritonX-100/PBS (PBT), and incubated overnight at 4 °C with the primary antibodies listed in Table 2. After 2 washes in PBT for 10 minutes each, the slides were incubated for 1 h at room temperature with the appropriate secondary antibodies listed in Table 2. Immunostained preparations were mounted in Vectashield H-200 with DAPI (Vector Laboratories). All cytological preparations were examined using either fluorescence microscope Nikon equipped with a Charged-Coupled Device (CCD camera; Photometrics CoolSnap HQ) or Zeiss LSM980 apparatus, using a 63x/1.40 NA oil objective.

**Table 2.**
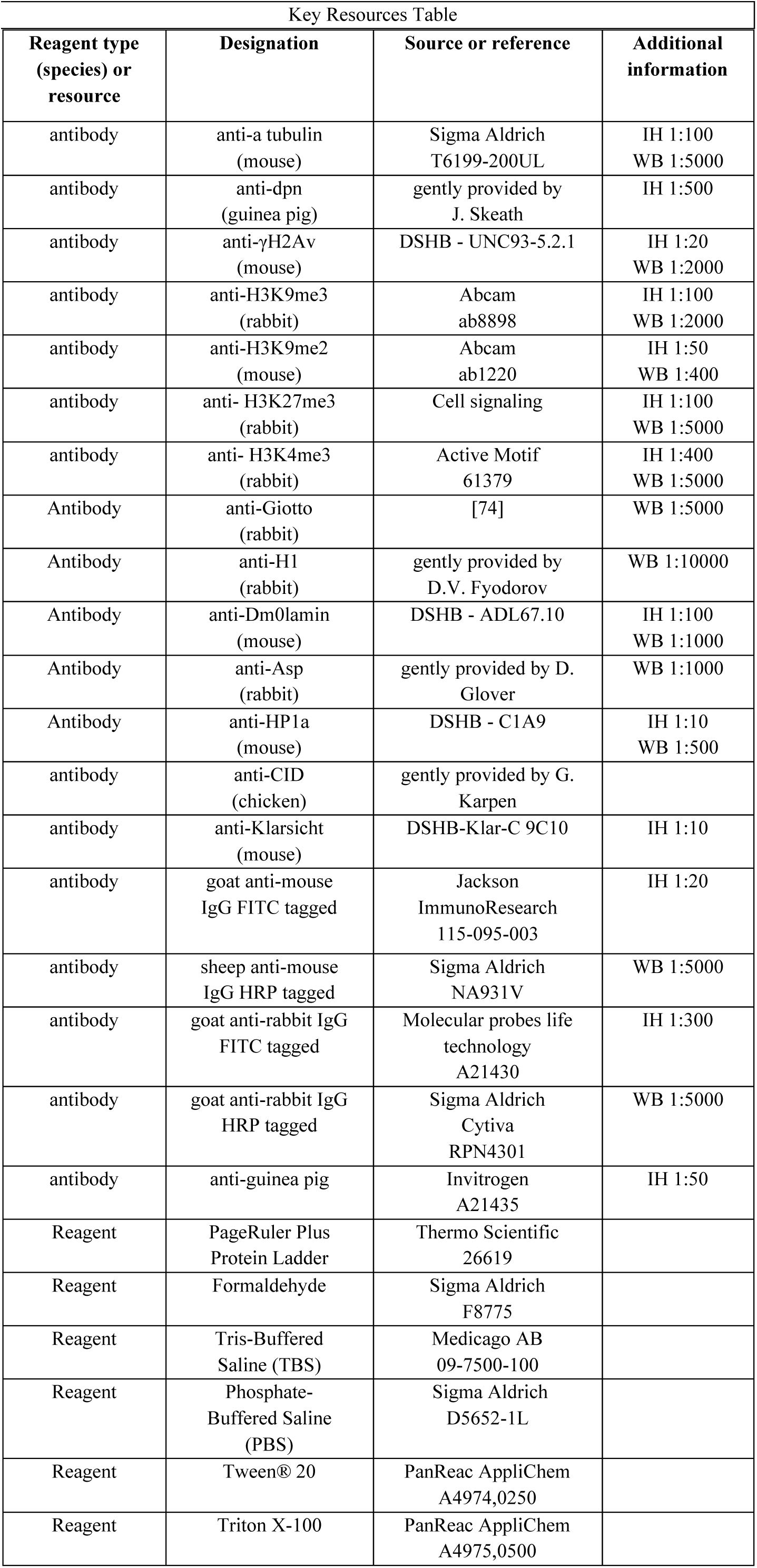
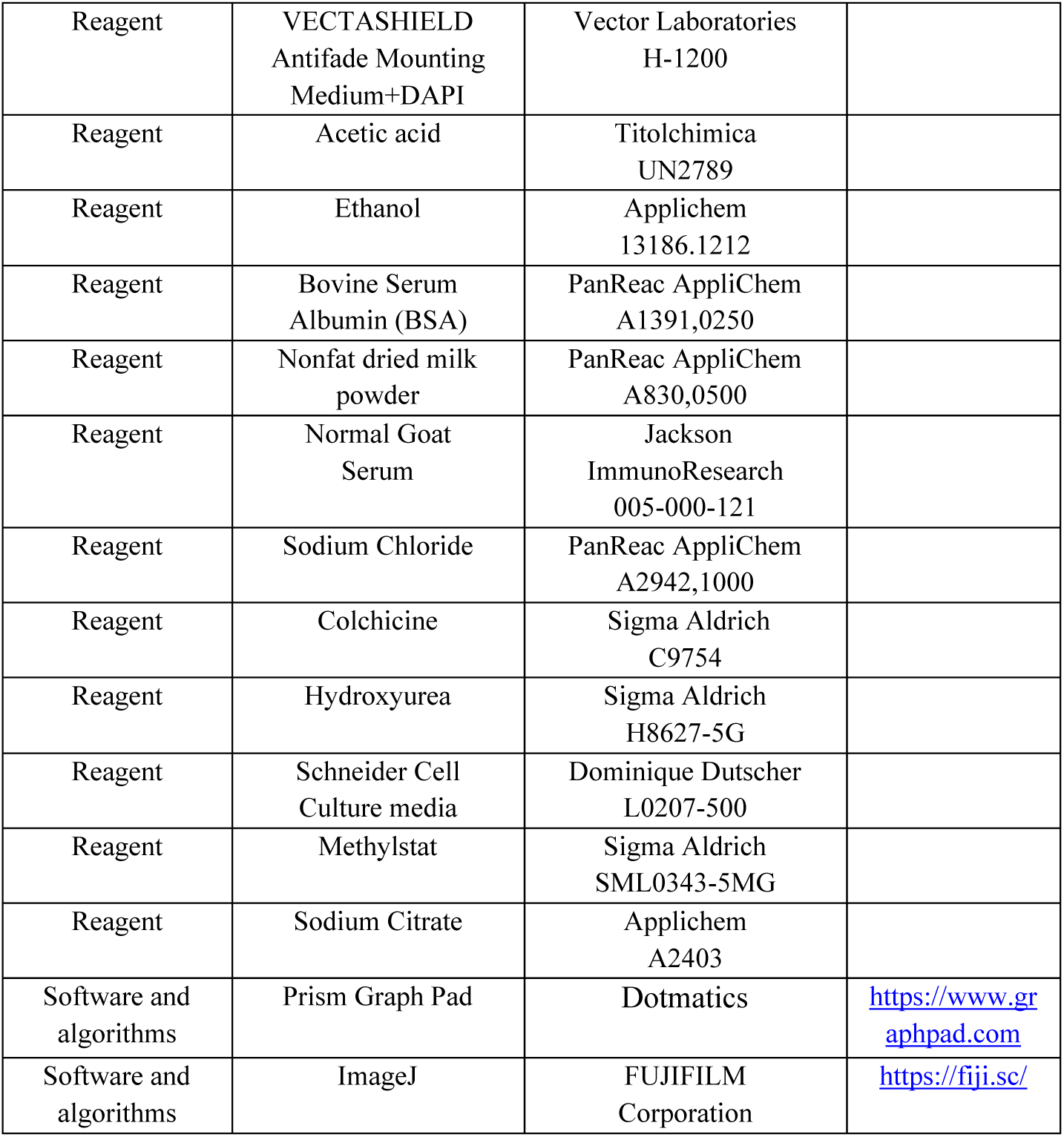

|  |
| --- |
| Key Resources Table |

| Reagent type<br>(species) or<br>resource | Designation | Source or reference | Additional<br>information |
| --- | --- | --- | --- |
| antibody | anti-a tubulin<br>(mouse) | Sigma Aldrich<br>T6199-200UL | IH 1:100<br>WB 1:5000 |
| antibody | anti-dpn<br>(guinea pig) | gently provided by<br>J. Skeath | IH 1:500 |
| antibody | anti-γH2Av<br>(mouse) | DSHB - UNC93-5.2.1 | IH 1:20<br>WB 1:2000 |
| antibody | anti-H3K9me3<br>(rabbit) | Abcam<br>ab8898 | IH 1:100<br>WB 1:2000 |
| antibody | anti-H3K9me2<br>(mouse) | Abcam<br>ab1220 | IH 1:50<br>WB 1:400 |
| antibody | anti- H3K27me3<br>(rabbit) | Cell signaling | IH 1:100<br>WB 1:5000 |
| antibody | anti- H3K4me3<br>(rabbit) | Active Motif<br>61379 | IH 1:400<br>WB 1:5000 |
| Antibody | anti-Giotto<br>(rabbit) | [74] | WB 1:5000 |
| Antibody | anti-H1<br>(rabbit) | gently provided by<br>D.V. Fyodorov | WB 1:10000 |
| Antibody | anti-Dm0lamin<br>(mouse) | DSHB - ADL67.10 | IH 1:100<br>WB 1:1000 |
| Antibody | anti-Asp<br>(rabbit) | gently provided by D.<br>Glover | WB 1:1000 |
| Antibody | anti-HP1a<br>(mouse) | DSHB - C1A9 | IH 1:10<br>WB 1:500 |
| antibody | anti-CID<br>(chicken) | gently provided by G.<br>Karpen |  |
| antibody | anti-Klarsicht<br>(mouse) | DSHB-Klar-C 9C10 | IH 1:10 |
| antibody | goat anti-mouse<br>IgG FITC tagged | Jackson<br>ImmunoResearch<br>115-095-003 | IH 1:20 |
| antibody | sheep anti-mouse<br>IgG HRP tagged | Sigma Aldrich<br>NA931V | WB 1:5000 |
| antibody | goat anti-rabbit IgG<br>FITC tagged | Molecular probes life<br>technology<br>A21430 | IH 1:300 |
| antibody | goat anti-rabbit IgG<br>HRP tagged | Sigma Aldrich<br>Cytiva<br>RPN4301 | WB 1:5000 |
| antibody | anti-guinea pig | Invitrogen<br>A21435 | IH 1:50 |
| Reagent | PageRuler Plus<br>Protein Ladder | Thermo Scientific<br>26619 |  |
| Reagent | Formaldehyde | Sigma Aldrich<br>F8775 |  |
| Reagent | Tris-Buffered<br>Saline (TBS) | Medicago AB<br>09-7500-100 |  |
| Reagent | Phosphate-Buffered Saline<br>(PBS) | Sigma Aldrich<br>D5652-1L |  |
| Reagent | Tween® 20 | PanReac AppliChem<br>A4974,0250 |  |
| Reagent | Triton X-100 | PanReac AppliChem<br>A4975,0500 |  |
| Reagent | VECTASHIELD<br>Antifade Mounting<br>Medium+DAPI | Vector Laboratories<br>H-1200 |  |
| Reagent | Acetic acid | Titolchimica<br>UN2789 |  |
| Reagent | Ethanol | Applichem<br>13186.1212 |  |
| Reagent | Bovine Serum<br>Albumin (BSA) | PanReac AppliChem<br>A1391,0250 |  |
| Reagent | Nonfat dried milk<br>powder | PanReac AppliChem<br>A830,0500 |  |
| Reagent | Normal Goat<br>Serum | Jackson<br>ImmunoResearch<br>005-000-121 |  |
| Reagent | Sodium Chloride | PanReac AppliChem<br>A2942,1000 |  |
| Reagent | Colchicine | Sigma Aldrich<br>C9754 |  |
| Reagent | Hydroxyurea | Sigma Aldrich<br>H8627-5G |  |
| Reagent | Schneider Cell<br>Culture media | Dominique Dutscher<br>L0207-500 |  |
| Reagent | Methylstat | Sigma Aldrich<br>SML0343-5MG |  |
| Reagent | Sodium Citrate | Applichem<br>A2403 |  |
| Software and<br>algorithms | Prism Graph Pad | Dotmatics | https://www.graphpad.com |
| Software and<br>algorithms | ImageJ | FUJIFILM<br>Corporation | https://fiji.sc/ |

### Polytene chromosome immunostaining

Salivary glands were dissected in physiological solution and fixed with 1.8% paraformaldehyde and 45% acetic acid for 7 minutes, then transferred onto a glass slide cover with a coverslip and then squahed with gentle tapping. The spread of polytene chromosome preparations was checked under microscope and then frozen in liquid nitrogen for 10 minutes. After removal of the coverslips, the slides were immersed in cold TBS for 10 minutes, then rinsed twice for 10 min each in 0.1% Tween-20/TBS (TBS-T) and then incubated with the appropriate primary antibodies listed in Table 2. The preparations were washed twice in TBS-T for 10 minutes each and incubated with the appropriate secondary antibodies, listed in Table 2, for 1 hour at room temperature. The preparations were then washed twice in TBS-T for 10 minutes each and mounted in Vectashield H-200 with DAPI (Vector Laboratories).

### Chromosome cytology

To analyze metaphase chromosomes, brains were dissected in physiological solution (0.7% NaCl), incubated with colchicine (10^-5^M in PBS) for 45 min, and then for 10min in hypotonic solution (0.5% Sodium Citrate). The preparations were squashed in 45 % acetic acid and frozen in liquid nitrogen for at least 10 minutes. The preparations were mounted in Vectashield H-200 with DAPI (Vector Laboratories).

### Immunoblot and antibodies

Protein extracts from *Drosophila* larval brains were obtained by dissecting 10 larval brains in 0.7% NaCl, diluted in 20 μl of 2X Laemmli buffer (4% SDS, 10% 2-mercaptoethanol, 20% glycerol, 0.004% bromophenol blue, 0.125 M Tris HCl, pH 6.8), boiled for 5 minutes and centrifugated at 7000 rpm for 30 seconds. Protein samples were loaded onto 15% SDS-PAGE gels and blotted onto PVDF or nitrocellulose filter membrane (Hybond ECL, Amersham). Filters were blocked in 5% non-fat dry milk dissolved in 0.1% Tween-20/PBS for 30 min at RT and then incubated with the appropriate antibodies, listed in Table 2, overnight at 4 °C. The blots were washed three times with 1X TBS-T for 5 minutes each, incubated with the appropriate HRP-conjugated secondary antibodies for 1 h at RT and then washed again 3 times with 0.1%Tween-20/PBS. The chemiluminescent signal was revealed through either SuperSignal™ West Femto or SuperSignal™ West Pico substrate (Thermo Scientific™) using the ChemiDoc scanning system (Bio-Rad).

### Quantification and statistical analysis

For analysis of the fluorescence intensity, the corrected total cell fluorescence (CTCF) was quantified using Image J software and calculated using the following formula: CTCF=Mean fluorescence intensity - (Area of selected nucleus x background mean fluorescence intensity). Western blot band intensity measurements were quantified by densitometric analysis using the Image Lab 4.0.1 software (Bio-Rad). WB was repeated independently at least three times.

Statistical analysis for comparison of two groups was done by t-test using GraphPad Prism 8.1.

## Acknowledgments

This work was supported by the National Recovery and Resilience Plan (NRRP), Mission 4, Component 2, Investment 1.1, Call for tender No. 104 published on 2.2.2022 by the Italian Ministry of University and Research (MUR), funded by the European Union – NextGenerationEU– Project 2022M75NN8, title ‘Dissection of common mechanisms in genetic primary microcephaly’. CUP B53D23008280006 and B53D23008270006 to LC and MPS, respectively. We thank Ludovica Picchetta for initial experiments, Maurizio Gatti and Ferdinando di Cunto for critical reading and scientific discussion, D.V. Fyodorov, G. Karpen and J. Skeath for generously providing us with antibodies and T. Schoborg for the *asp^t25^* stock. We also thank the Bloomington Stock Center for the fly stocks.

## Legends

**Supplementary figure 1:**
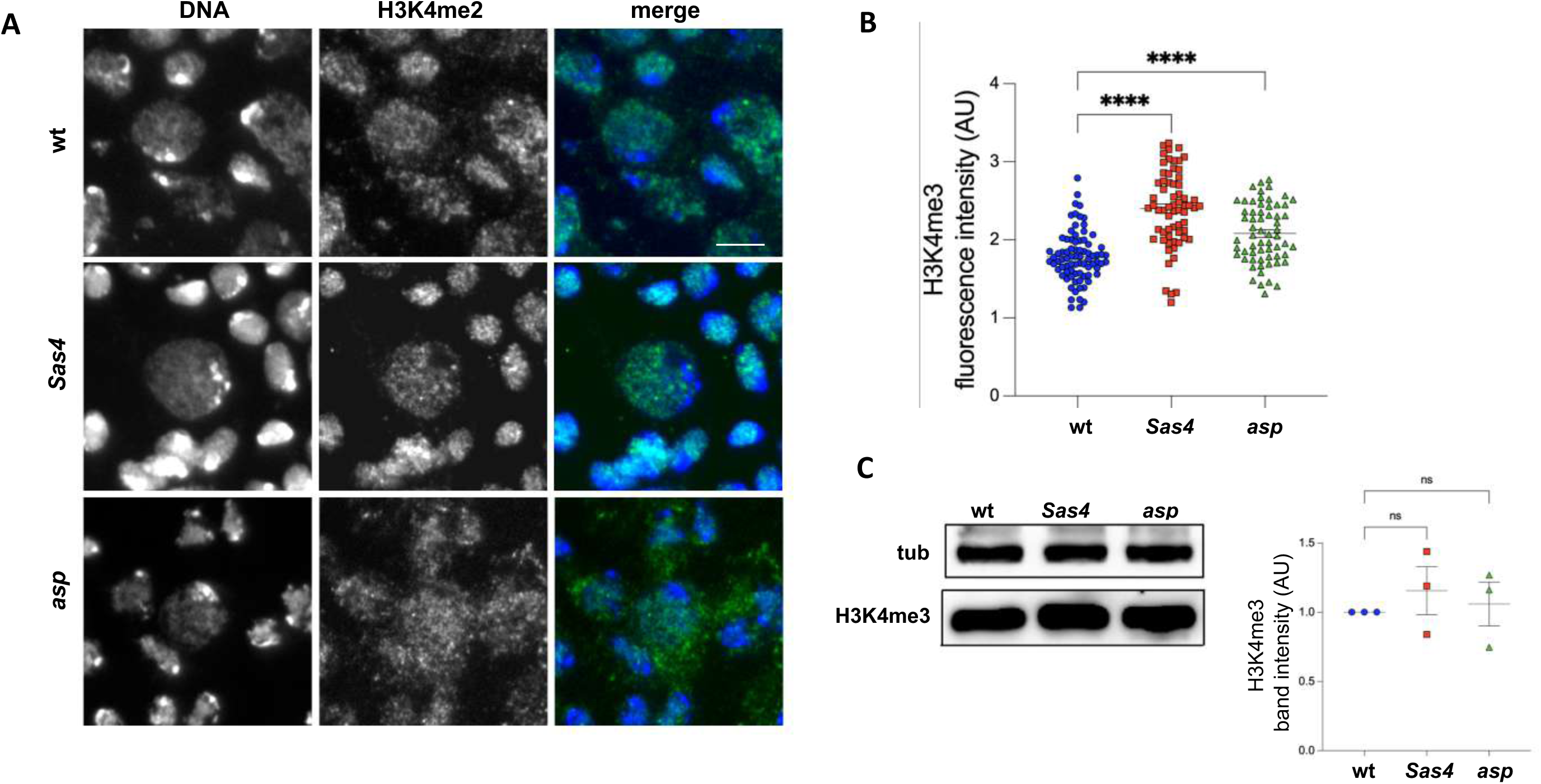
Loss of Sas4 or Asp alters H3K4me3 levels in neuroblasts. A) Immunolocalization of H3K4me3 on brain squashes from third instar larvae of wt and *Sas4* or *asp* mutants. In the merged panels, H3K4me3 is in green and DNA in blue (DAPI). Scale bar = 10 μm. B) H3K4me3 fluorescence intensity quantification in neuroblasts, showing an increase of H3K4me3 in both *Sas4* and *asp* mutants compared to wt, each dot represents a single cell (n ≥50). C) Representative western blots showing no change of H3K4me3 in larval brain extracts of *Sas4* or *asp* mutants in comparison to wt extracts with the corresponding band quantification normalized on the loading control (Tubulin). AU, arbitrary unit. Error bars represent SEM. P = p-value calculated using unpaired t test. *p < 0.05; **p < 0.01; ***p < 0.001; ****p < 0.0001.

**Supplementary figure 2:**
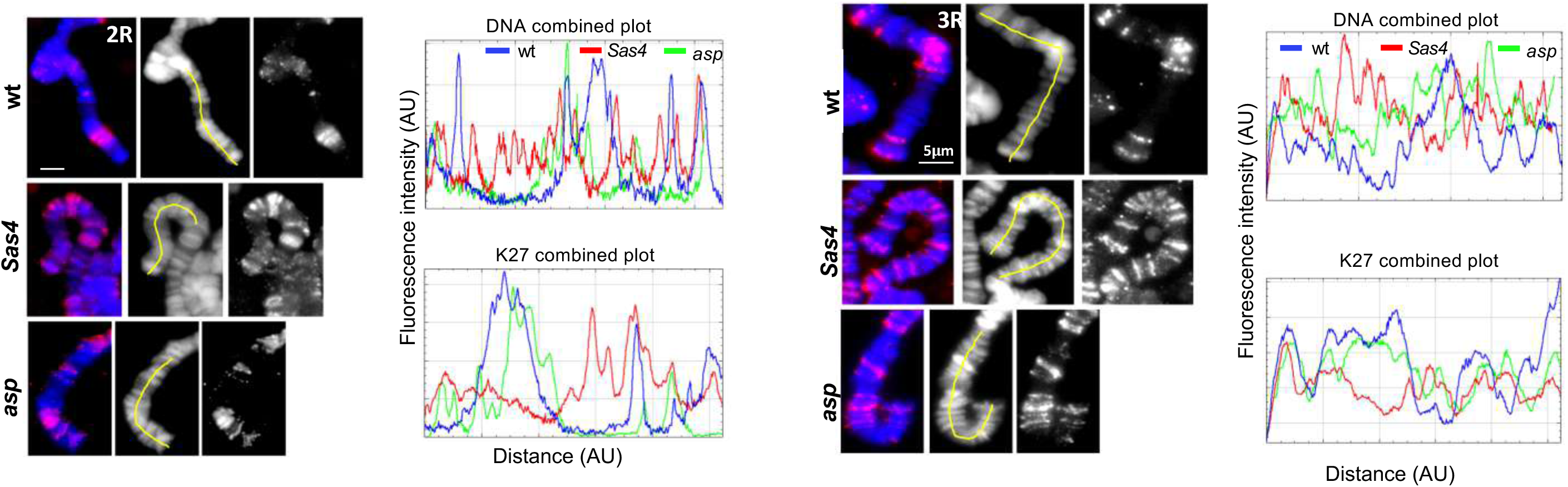
DNA and H3K27me3 banding patterns are altered in *Sas4* and *asp* mutant polytene chromosomes. Examples of the 2R and 3R chromosome extremities stained with anti H3K27me3 (red) and DNA (blue) in wt and *Sas4* or *asp* mutant polytene chromosomes with corresponding intensity profiles, showing an altered pattern of DNA and H3K27me3 signals in both *Sas4* (red line) and *asp* (green line) mutants compared to wt (blue line); y-axis: fluorescence intensity; x-axis: distance from the tip of the chromosome (arbitrary unit). Scale bar = 5 μm.

**Supplementary figure 3:**
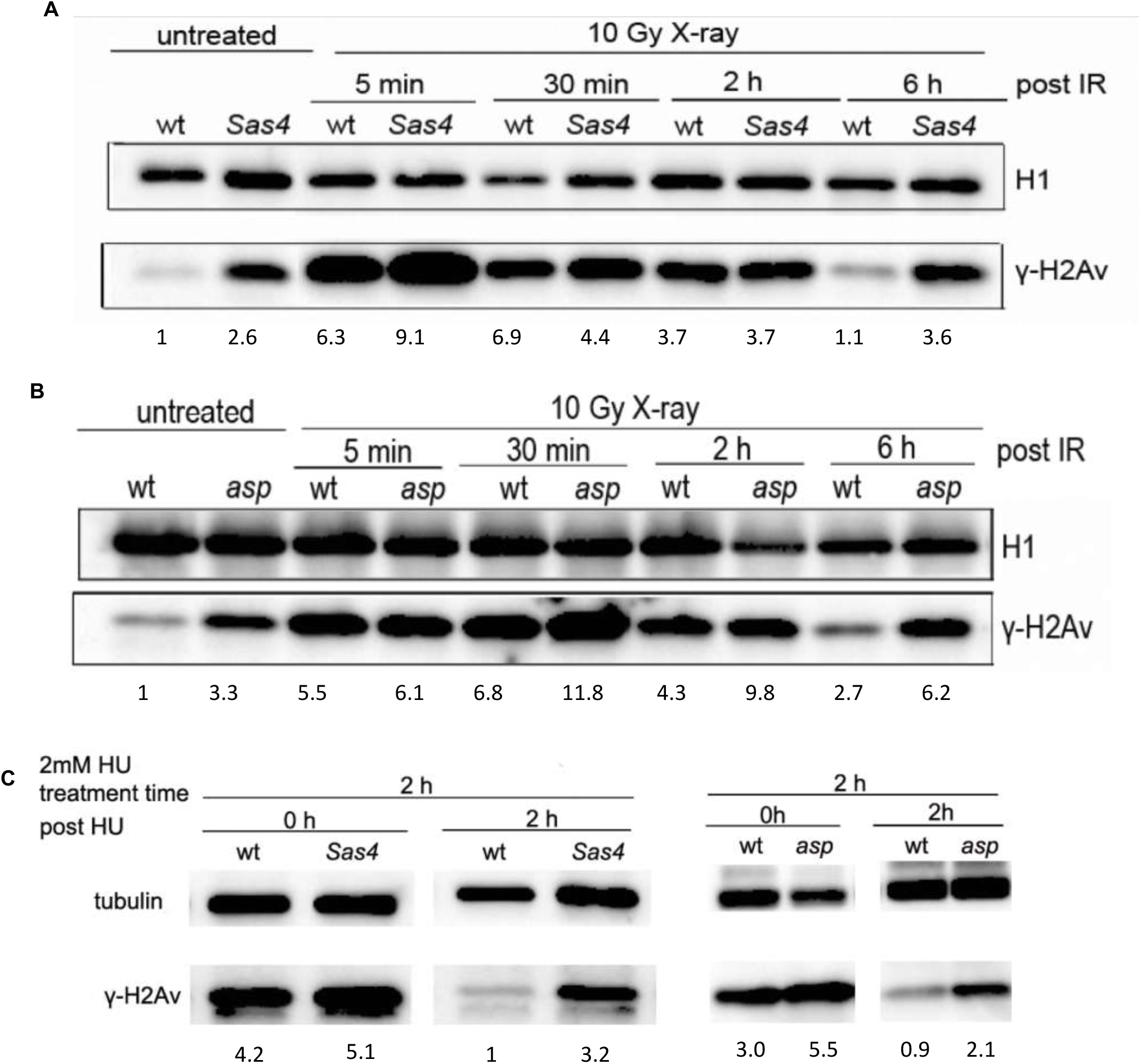
*Sas4* and *asp* mutant brain cells exhibit delayed DNA Damage Response following irradiation and hydroxyurea treatment. A) γH2Av levels in wt and *Sas4* larval brain extracts after irradiation with 10Gy and dissection after 5 min, 30 min, 2 hours and 6 hours of recovery time. H1 was used as loading control and the γH2Av band quantification was normalized on the loading control. B) γH2Av levels in wt and *asp* larval brain extracts after irradiation with 10Gy and dissection after 5 min, 30 min, 2 hours and 6 hours of recovery time. H1was used as loading control and the γH2Av band quantification was normalized on the loading control. C) γH2Av levels in wt or *Sas4* and *asp* mutant larval brain extracts detected with γH2Av antibody and tubulin as loading control. Larval brains were treated with 2mM of HU for 2 hours and dissected at 0 and 2 hours post treatment (recovery time). The γH2Av band quantification was normalized on the loading control.

**Supplementary figure 4:**
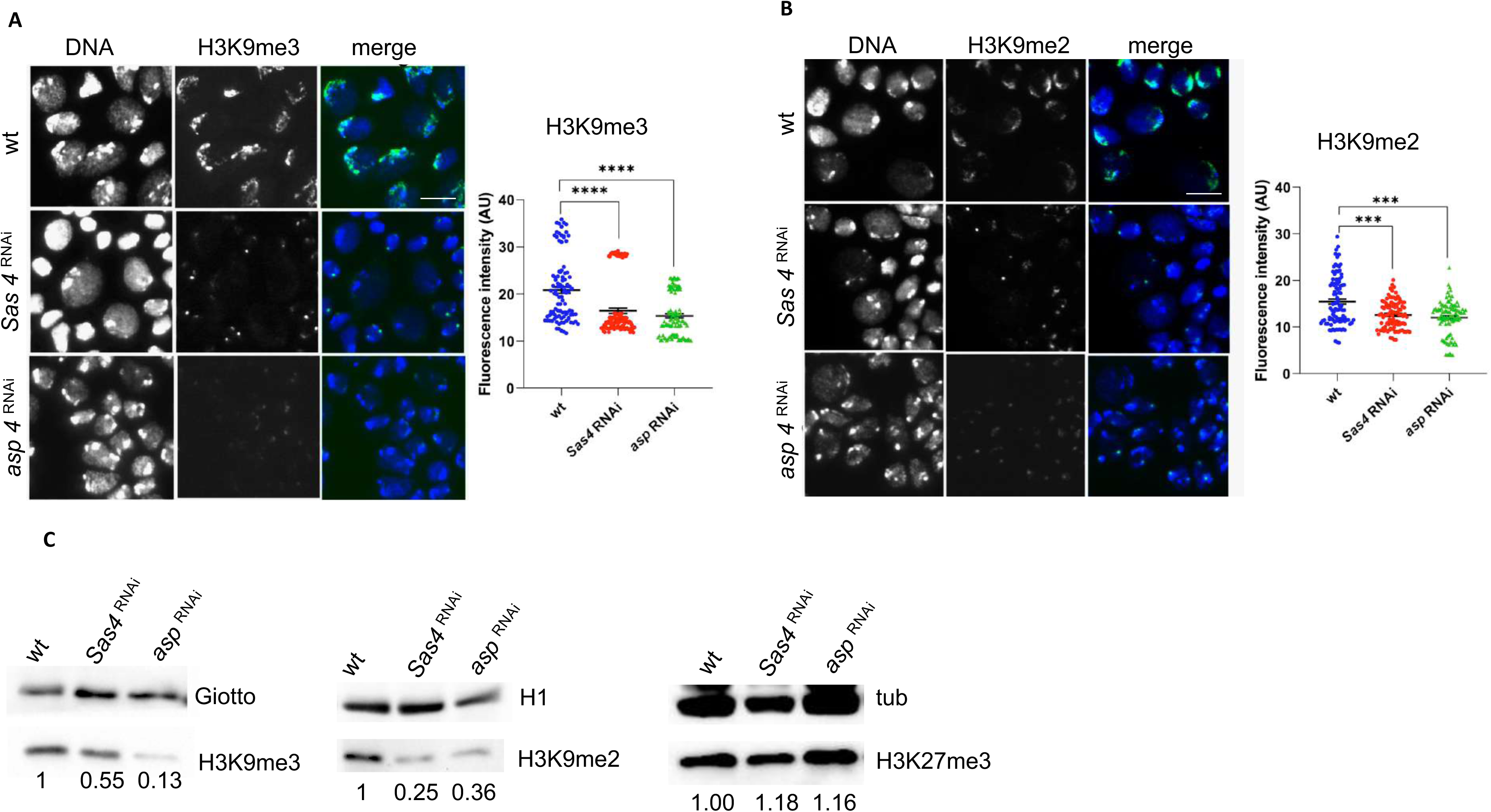
RNAi-mediated silencing of *Sas4* or *asp* recapitulates the heterochromatin marks reduction of mutant phenotypes. A) Third instar brain squashes of wt and *Tub-GAL4*>*UAS-Sas4^RNAi^* or *UAS-asp^RNAi^* immunostained with anti-H3K9me3 and quantification of H3K9me3 fluorescence intensity per cell as in (A), each dot represents a single cell from at least three brains (n ≥ 50). Scale bar = 10 μm. B) Third instar brain squashes of wt and *Tub-GAL4*>*UAS-Sas4^RNAi^* or *UAS-asp^RNAi^* immunostained with anti-H3K9me2 and quantification of H3K9me2 fluorescence intensity per cell as in (B), each dot represents a single cell from at least three brains (n ≥ 50). C) Immunoblots showing reduced levels of H3K9me3 and me2 and no difference in H3K27me3 level upon *Sas4* or *asp* silencing compared to wt. The corresponding band quantification was normalized on the loading control (Giotto, H1 or tubulin).

**Supplementary figure 5:**
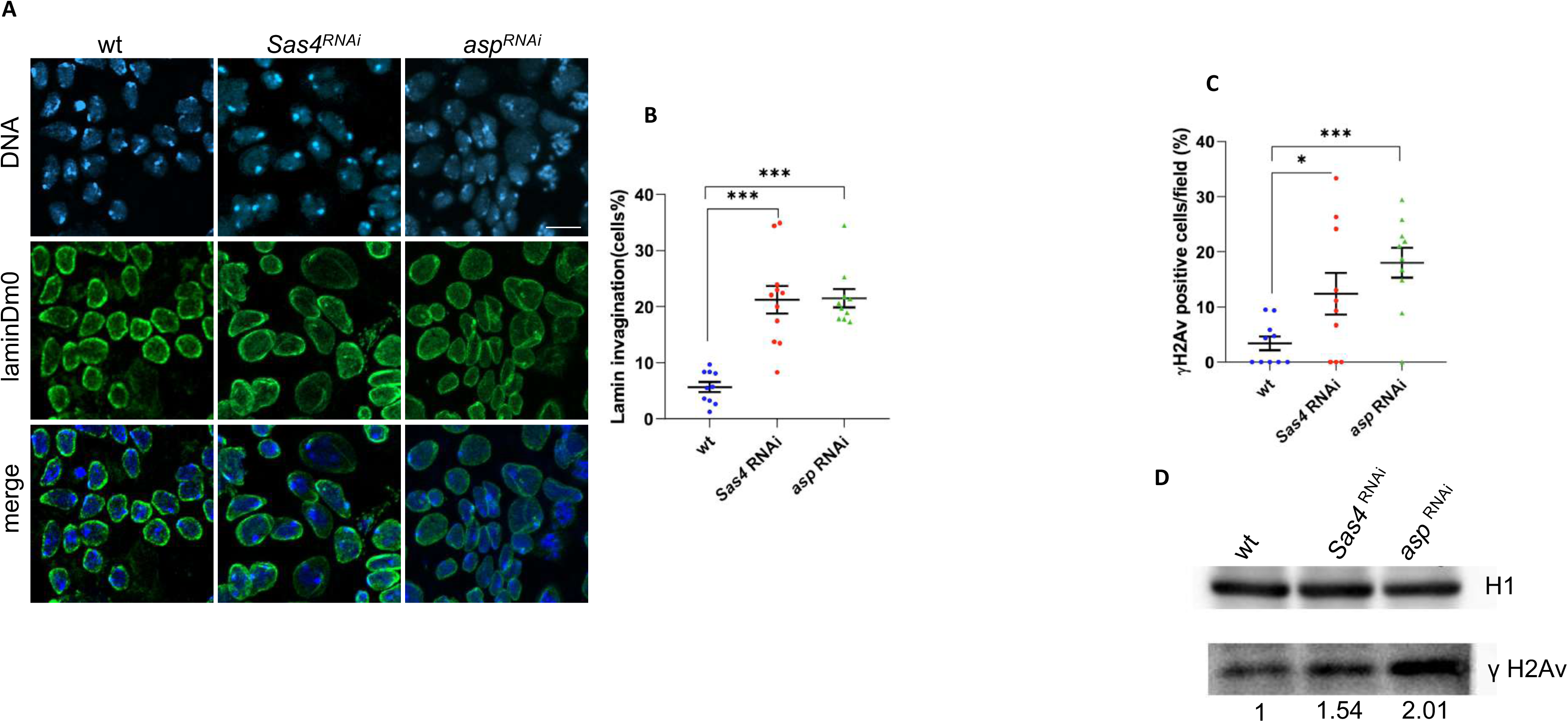
RNAi-mediated silencing of *Sas4* or *asp* recapitulates both the lamin alteration and the increased levels of γH2Av of mutant phenotypes. A) Third instar brain squashes of wt and *Tub-GAL4*>*UAS-Sas4^RNAi^* or *UAS-asp^RNAi^* immunostained with anti-LaminDM0 antibody (green) and DAPI (blue, DNA). Note that all cell types exhibit lamin invaginations. Scale bar = 10 μm. B) Graphical representation of the percentage of cells showing invaginations of the nuclear envelope in *Tub-GAL4*>*UAS-Sas4^RNAi^*(red rectangles) and *UAS-asp^RNAi^* (green triangles) and wt (blue circles) brain cells. Each dot represents the score of cells per 63x microscope field (n≥10). C) Graphical representation of the percentage of γH2Av positive cells per field, n≥10. D) Immunoblots on brain extracts from wt and *Tub-GAL4*>*UAS-Sas4^RNAi^* or *UAS-asp^RNAi^* labelled using anti-γH2Av antibody with the corresponding band quantification normalized on the loading control (H1). Error bars represent SEM. P = p-value calculated using unpaired t test. *p < 0.05; **p < 0.01; ***p < 0.001; ****p < 0.0001.

